# C9ORF72-derived polyGR polypeptides disrupt passive nucleocytoplasmic transport by tuning protein affinity for the nuclear pore barrier

**DOI:** 10.64898/2026.03.16.711670

**Authors:** Daniel A Solomon, Ryan J Emenecker, Marie-Therese Salcher-Konrad, Sofia Marina Konstantinidou, Olivia H Houghton, Eleanor Wycherley, Seoungjun Lee, Niamh L O’Brien, Juan Alcalde, Inês Lourenço Cabaço, Marc-David Ruepp, H Broder Schmidt, Alex S Holehouse, Sarah Mizielinska

**Author notes:** These authors contributed equally to this work (Daniel A Solomon, Ryan J Emenecker, Marie-Therese Salcher-Konrad).

## Abstract

Nucleocytoplasmic partitioning is an essential determinant of eukaryotic cellular function, governed by the nuclear pore complex, a molecular portal filled by a disordered phenylalanine-glycine (FG)-rich phase that governs selective entry and exit. Disruption of nucleocytoplasmic partitioning is seen across many different diseases, including viral infection, cancer, and neurodegeneration. However, what determines whether a given protein mislocalises when nucleocytoplasmic transport is disrupted remains unknown. This question is central to amyotrophic lateral sclerosis and frontotemporal dementia (ALS/FTD), where cytosolic mislocalisation of the nuclear RNA-binding protein TDP-43 is a defining pathological feature. The most common genetic cause of these diseases is a G_4_C_2_ repeat expansion in the gene *C9ORF72*, which produces aberrant neurotoxic polypeptides that induce nucleocytoplasmic transport defects. Here, we show how the highly toxic poly(glycine-arginine/GR) polypeptide engages the nuclear pore FG-rich selectivity barrier and retunes passive nucleocytoplasmic transport according to client surface chemistry. Using coarse-grained simulations, *in vitro* FG-phase reconstitution and human cell lines and neurons, we find that polyGR produces a non-linear, biphasic modulation of nuclear pore passage. Proteins with low affinity for the FG phase are unaffected, whereas proteins with higher affinity due to solvent-exposed hydrophobic residues exhibit enhanced transport up to a critical threshold, beyond which highly hydrophobic proteins experience transport suppression, cytoplasmic accumulation and aggregation. Together, these findings establish how disease-associated polypeptides retune the physicochemical rules governing passive nuclear pore transport, leading to biphasic outcomes determined by protein surface chemistry that alter protein compartmentalisation and aggregation. This provides a biophysical mechanism by which polyGR drives selective protein vulnerability to nuclear pore dysfunction in *C9ORF72*-associated ALS/FTD.

## Introduction

Nuclear compartmentalization is a key factor in the regulation of gene expression by spatial segregation, critical for many aspects of cellular homeostasis and response to environmental cues or stressors. Transport between the nucleus and cytoplasm, termed nucleocytoplasmic transport, is tightly regulated by nuclear pore complexes, which serve as gateways controlling the import and export of proteins, RNAs, and other macromolecules^1, 2^. The nuclear pore complex (NPC) allows two modes of transport: passive diffusion and active translocation^2^. At the heart of both is a permeability barrier formed by phenylalanine-glycine nucleoporins (FG Nups)^1, 2^. In active nucleocytoplasmic transport, cargoes bearing nuclear localisation or export signals are shuttled across the FG Nup barrier by nuclear transport receptors in a process driven by the Ran-GTPase cycle^3^. In passive diffusion, molecules typically smaller than 30-40 kDa pass unassisted through the FG Nup barrier^2^.

Disruption of the compartmentalization of nuclear factors critical for gene expression is a hallmark of several neurodegenerative disorders, particularly amyotrophic lateral sclerosis (ALS) and frontotemporal dementia (FTD)^4–8^. For example, nuclear loss and cytoplasmic accumulation of the DNA/RNA-binding protein TDP-43 is a hallmark pathology in ALS and FTD, present in ∼97% and ∼45% of cases, respectively^9^. A hexanucleotide repeat expansion mutation in the gene *C9ORF72* is the most common genetic cause of ALS and FTD and is also associated with TDP-43 pathology^9–13^. This mutation results in the production of aberrant polypeptides^14, 15^, of which the two arginine-rich polypeptides polyGR and polyPR are the most toxic^16–18^. Defective transport through the NPC has emerged as a key pathogenic pathway in ALS/FTD^4, 5, 8, 19–21^ and is also associated with the *C9ORF72* arginine-rich polypeptides^19, 22–27^. Understanding the mechanisms underlying these deficits is critical for understanding disease progression and the cellular basis of neurodegeneration^4^.

Interestingly, the *C9ORF72* arginine-rich DPRs preferentially bind to proteins with low complexity domains^28, 29^. These domains, which are often intrinsically disordered^30^, enable proteins to undergo multivalent interactions with other proteins or nucleic acids^31–33^, a process that underlies phase separation and often biomolecular condensate formation^31, 34, 35^. The permeability barrier of the NPC is formed by intrinsically disordered, low-complexity FG-nucleoporins (e.g. Nup98), whose FG-repeat domains engage in dynamic multivalent interactions that create a selective “FG phase” within the central channel^1, 2^. An unresolved question is whether nucleocytoplasmic transport disruptions observed in *C9ORF72*-ALS/FTD are associated with pathological interactions between arginine-rich DPRs and the low complexity domains of FG Nups.

Using a combination of cellular models, *in vitro* assays, and coarse-grained simulations, we show that the aberrant *C9ORF72* polypeptide polyGR partitions into condensates formed by the FG domain of the human nucleoporin Nup98, thereby altering how proteins engage the FG phase and traverse the nuclear pore. Strikingly, these effects are primarily dictated by client protein surface chemistry, producing biphasic outcomes spanning enhanced nucleocytoplasmic transport, transport suppression, and protein aggregation. Together, these findings uncover a mechanism by which polyGR selectively reprograms passive diffusion through the nuclear pore in a protein-specific manner, providing key mechanistic insight into nucleocytoplasmic transport defects that contribute to *C9ORF72*-associated ALS and FTD, and more generally explaining protein vulnerability to NPC dysfunction.

## Results

### Transport through the nuclear pore is altered by the aberrant peptide polyGR in a protein-specific manner

Passive transport, the diffusion of client molecules across the NPC, is primarily governed by cohesive FG–FG interactions among FG Nups^2, 36–38^. Here, we use the term “client” to refer to any molecular species that interacts with or partitions into the nuclear pore FG phase, including passive proteins and transport receptors; the term “cargo” is reserved for species recognised and carried by nuclear transport receptors^31, 39^. Passive transport of a given client across the NPC is dependent on its size; without favourable FG domain interactions, molecules >30 kDa diffuse poorly, whereas smaller molecules cross rapidly^2, 40^. The NPC therefore acts – at least in part – as a chemically tuned kinetic barrier, with transit times increasing with client size and FG-interaction propensity^2, 41^. As passive transport is mechanistically independent of transport receptors and the Ran gradient^42^, monitoring this process is a direct readout of NPC integrity and can reveal FG Nup-related barrier defects in disease.

To address the impact of the *C9ORF72* neurotoxic polypeptide polyGR, we studied passive transport in cells upon treatment with a 40-mer of GR (GR_20_), herein referred to as polyGR. We first examined the effect of polyGR on size-dependent passive transport using dextrans of different molecular weights as clients. Employing a classical cell permeabilization nuclear import assay^43, 44^ (Extended Data Fig. 1a), we assessed passive import of fluorescently labelled dextrans (20, 40, 70 kDa) following treatment with polyGR, and two other non-toxic *C9ORF72* polypeptides polyGP and polyPA in HeLa cells using live-cell imaging. The 20 kDa dextran readily entered nuclei and accumulated in nucleolar-like structures, with polyGR causing a marked increase in transport compared to untreated cells (Fig. 1a,b; Extended Data Fig. 1b), as seen in a previous study^26^. In contrast, neither polyGP nor polyPA altered transport (Fig. 1a,b; Extended Data Fig. 1b). PolyGR exerted a milder increase in import for the 40 kDa dextran, which was again not seen with the polyGP control (Extended Data Fig. 1c,d), and finally, polyGR had no impact on the partitioning of the 70 kDa dextran (Extended Data Fig. 1e,c-f). Thus, polyGR disrupts passive nucleocytoplasmic transport in cells, with a greater impact on smaller clients.

**Figure 1.**
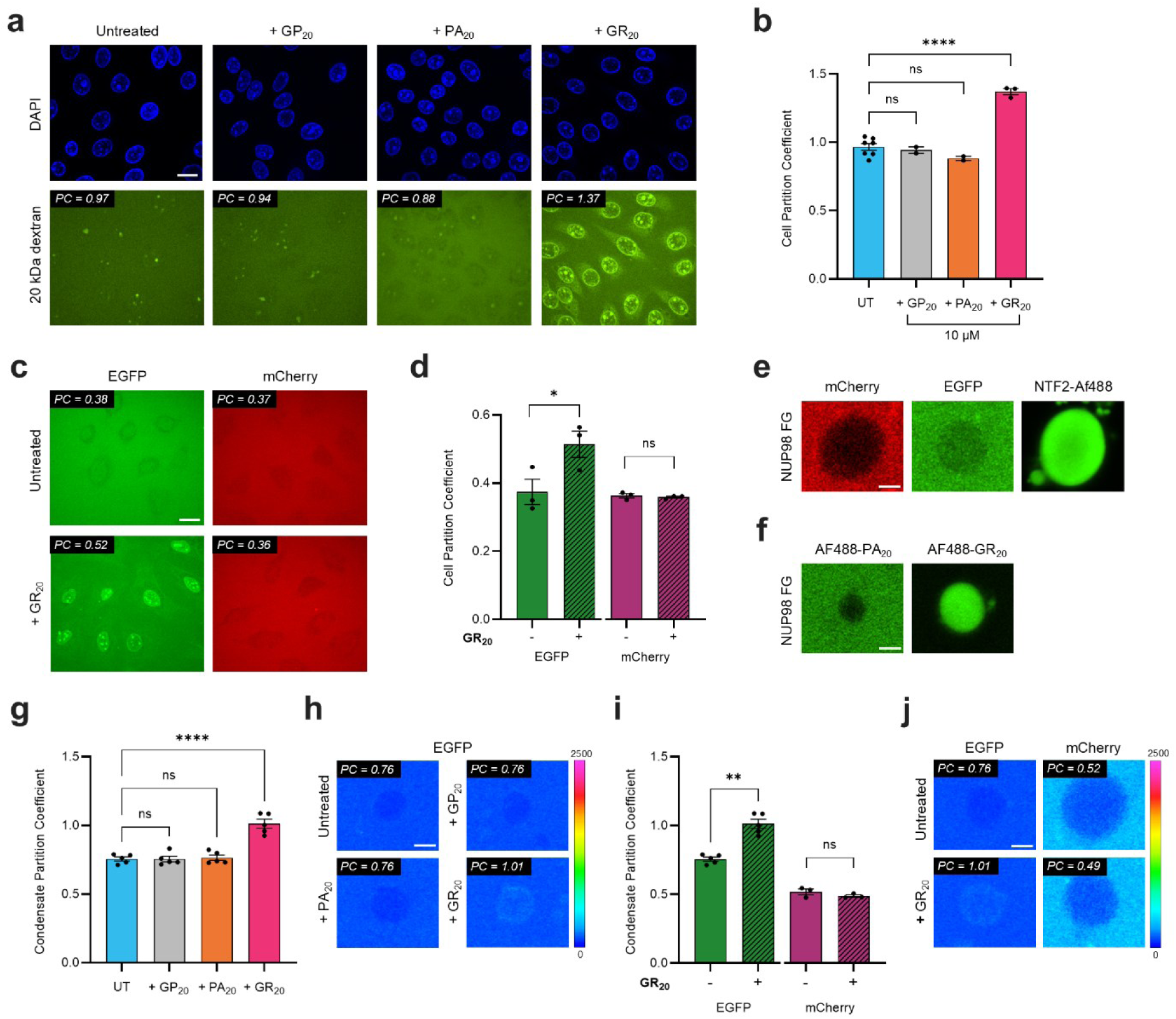
PolyGR binds nuclear pore FG phases and selectively enhances passive nuclear transport in a client-specific manner. **a,b,** Passive nuclear import assays in permeabilized HeLa cells reveal that 10 µM GR_20_ peptide markedly increases nuclear accumulation of 20 kDa fluorescent dextran compared with untreated (UT), GP_20_, or PA_20_ – representative images and mean cellular partition coefficient (nucleoplasm to background intensity ratio). Stats: one-way ANOVA with Tukey’s multiple comparisons test; n= 7 biological replicates for untreated, 3 for polyGR, 2 for polyGP and polyPA. **c,d,** GR_20_ selectively enhances nuclear import of EGFP but not mCherry (experiments performed separately but shown on same graph for ease of comparison). Stats: two-tailed paired *t*-test; n= 3 biological replicates. **e,** Recombinantly purified human Nup98 FG domain forms condensates that model NPC permeability; inert clients mCherry and EGFP are excluded from the Nup98 FG phase, whereas the transport receptor NTF2 readily partitions into the Nup98 FG phase. **f,** Fluorescently (Alexa488)-labelled GR_20_, but not PA_20_, partitions into Nup98 FG condensates, demonstrating direct and specific engagement with the FG network. **g,h**, Consistent with cellular data, GR_20_ increases EGFP partitioning into Nup98 FG condensates compared to untreated, whereas GP_20_ and PA_20_ have no effect; images show representative condensate intensity with heatmap scaling, together with corresponding mean condensate partition coefficients (PC) (inside to outside particle intensity ratio). Stats: one-way ANOVA with Tukey’s multiple comparisons test; n = 5 biological replicates, ≥5 condensates per replicate. **i,j** Consistent with cellular data, GR_20_ does not alter mCherry partitioning into Nup98 FG condensates relative to untreated unlike EGFP (replotted from Fig. 1g,h for ease of comparison); images show representative condensate intensity with heatmap scaling, together with corresponding mean condensate partition coefficients. Stats: two-tailed paired *t*-test: n= 5 biological replicates for EGFP, 3 for mCherry, ≥5 condensates per replicate. Data are shown as mean ± SEM. \*\*\*\**p* < 0.0001; \*\**p*<0.01; \**p*<0.05; ns, not significant. Scale bars = 20 µm for cells, 2 µm for condensates.

While dextran clients revealed a modulation of NPC permeability by polyGR, dextrans are non-physiological polymers and may not represent effects on proteins. To explore this, we examined the proteins EGFP and mCherry. Strikingly, polyGR enhanced nuclear import of EGFP (partition coefficient change, ΔPC = +0.14) while mCherry was unaffected (ΔPC = - 0.01) (Fig. 1c, d). This is surprising since EGFP and mCherry are approximately the same molecular weight (∼27 and ∼29 kDa, respectively) and share a highly similar beta-barrel structure. This suggests that, beyond molecular weight, intrinsic, protein-specific features determine susceptibility to polyGR-mediated modulation of passive nucleocytoplasmic transport.

### PolyGR engages nuclear pore FG domains via aromatic and hydrophobic interactions to alter barrier selectivity

To directly test how molecules interact with the NPC barrier, we turned to a reductionist system using *in vitro* FG-rich condensates^37^. Although these condensates do not reproduce the full nanoscale architecture of intact NPCs^45^ they provide a valuable and widely used model to study FG molecular interactions^2, 36–38, 41, 46–52^.

Several FG domains of FG-Nups phase separate into condensates *in vitro*^37, 49, 53^ and exhibit NPC-like transport selectivity, thereby permitting the partitioning of nuclear transport receptors and their cargoes while restricting inert molecules^37, 49^. We focused on the FG domain of human Nup98, a critical component of the vertebrate permeability barrier^38^. The Nup98 FG domain accounts for the largest FG mass in the NPC, and the transport selectivity of Nup98 FG condensates has been well-characterised across species^36, 37, 39, 41, 50, 54–57^. Further, Nup98 is strongly linked to several human neurodegenerative diseases^58–63^.

We purified the highly disordered FG domain of human Nup98 (amino acids 1-499) under denaturing conditions due to its high phase separation propensity (Extended Data Fig. 2a,b). Upon dilution of the denaturant, the Nup98 FG domain spontaneously phase-separates into condensates (referred to herein as FG condensates) (Extended Data Fig. 2c). We confirmed NPC-like functional selectivity of our Nup98 FG condensates, demonstrating that they exclude inert molecules like EGFP and mCherry, whilst allowing selective partitioning of Alexa488-labeled nuclear transport receptor nuclear transport factor 2 (NTF2) (Fig. 1e). Thus, our recombinantly purified human Nup98 FG condensates recapitulate the selective partitioning behaviour characteristic of FG-mediated NPC transport, consistent with their long-standing use as biochemically faithful models for probing interactions between the FG phase and its different clients ^36, 37, 39, 41, 50, 56, 57^.

Prior work has established that the *C9ORF72* polypeptide polyGR is a promiscuous interactor and partitions into various condensates^17, 28, 64–69^. We therefore examined polyGR interactions with the human Nup98 FG phase. Fluorescently labelled polyGR readily partitions into Nup98 FG condensates, whereas polyPA does not, demonstrating specific molecular requirements for interaction (Fig. 1f). To elucidate these requirements, we created Nup98 FG variants: i) F>Y (phenylalanine to tyrosine) – to preserve aromatic interactions and hydrophobicity, ii) F>L (phenylalanine to leucine) – to retain hydrophobic motifs but eliminate aromatic interactions, and iii) F>S (phenylalanine to serine) – to eliminate aromatic interactions and hydrophobic motifs^47^ (Extended Data Fig. 2c). The F>Y variant, like the wild-type FG domain, readily formed condensates upon dilution; in contrast, at the same concentration, the F>L variant required an unphysiologically-high concentration of molecular crowder (80 mg/ml 75 kDa Dextran); finally, the F>S variant exhibited significantly reduced phase separation propensity, only forming a small number of condensates with an even higher concentration molecular crowder (150 mg/ml 75 kDa Dextran) (Extended Data Fig. 2d). PolyPA was fully excluded from the F>Y condensates, but not as strongly from the F>L and F>S condensates, indicative of a slight increase in condensate permeability (Extended Data Fig. 2f). PolyGR partitioning was strongest for wild-type FG and F>Y condensates, intermediate in F>L, and weakest in the F>S variant (Extended Data Fig. 2e). Although this does not rule out a contribution from other interactions, these results suggest that polyGR primarily engages with nuclear pore FG domains through aromatic interactions, with additional contributions from hydrophobicity.

We next examined how polyGR influences the selectivity of the human Nup98 FG phase. Surprisingly, while EGFP was excluded from FG condensates in the absence of polyGR, the presence of polyGR enhanced EGFP entry (Fig. 1g,h). This effect was not observed for polyPA or polyGP. Further, the impact of polyGR on EGFP partitioning was dependent on both peptide length and concentration; higher concentrations (Extended Data Fig. 3a,b) and longer peptides (Extended Data Fig. 3c,d) resulted in greater increases in EGFP partitioning. To determine whether enhanced influx was due to changes in permeation client dynamics in the FG phase, we performed fluorescence recovery after photobleaching (FRAP) on the nuclear transport receptor NTF2 partitioned into Nup98 FG condensates. However, FRAP revealed no change in NTF2 mobility in the presence of polyGR (Extended Data Fig. 3e,f), indicating that polyGR does not result in more dynamic NTF2 interactions in general. To test the selectivity of polyGR, we examined the influx of mCherry. As per the cell nuclear import experiments, despite having a similar molecular weight and globular structure to EGFP, polyGR did not enhance mCherry partitioning into Nup98 FG condensates (Fig. 1i,j). This protein-specific effect, consistent between isolated condensates and an intact cellular system, suggests that the disease-associated *C9ORF72* polyGR peptide can selectively modulate nuclear pore permeability depending on the physicochemical properties of the client protein.

### Simulations reveal the amino acid level impact of aberrant polyGR peptide on FG phase partitioning

Our work thus far suggests that the polyGR-dependent recruitment of a client into an FG phase is influenced – or even determined – by the surface chemistry of that client. To examine the physicochemical determinants of nuclear pore interactions, we turned to coarse-grained simulations. Consistent with our *in vitro* results, simulations using a representative fragment of Nup98 FG (Nup98^2-151^) formed a single condensate. To systematically evaluate NPC FG domain-peptide interactions, we simulated every possible combination of amino acid dipeptide repeat as a 10-mer in the presence of the Nup98 FG condensate, yielding a total of 210 distinct combinations. Further, we carried out these simulations in the presence or absence of Nup98-stoichiometrically matched polyGR 20-mer (Fig. 2a), reasoning that the dipeptide 10-mers would act as ‘chemical probes’ to assess the emergent recruitment or exclusion of different sets of amino acids into Nup98 FG condensates upon addition of polyGR.

**Figure 2.**
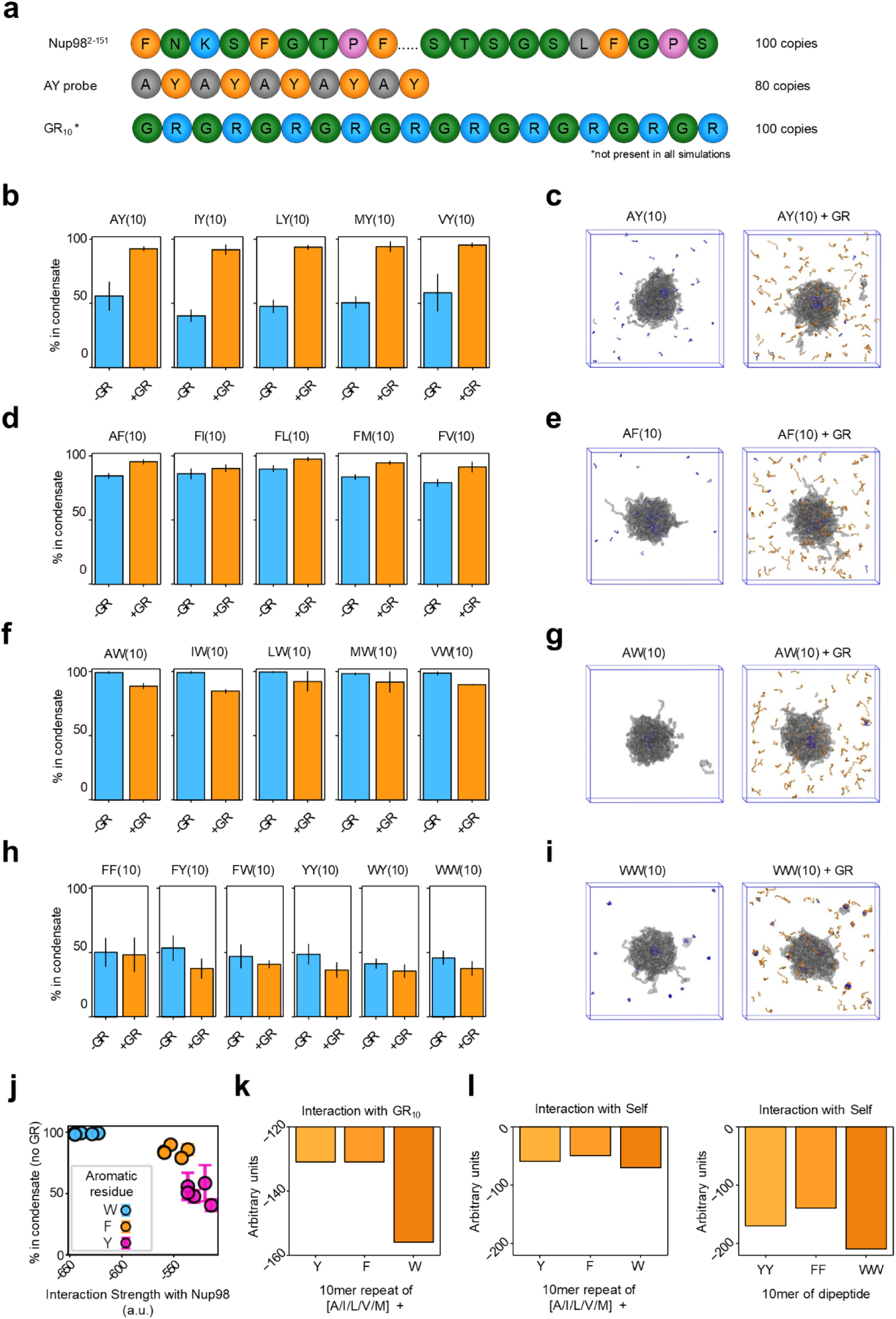
Coarse-grained simulations define the amino acid selectivity of polyGR-induced alterations in molecular interaction with the Nup98 FG phase. **a,** Schematic of coarse-grained simulation setup using equilibrated Nup98 FG domain (amino acids 2-151) condensates with 80 copies of 10-mer dipeptide repeat probes to assess amino acid selectivity and, where indicated, 100 copies of GR_10_ polypeptide. **b,d,f,h,** Bar graphs show the average percentage of each indicated probe that is inside of the Nup98 FG domain condensate from two independent replicates of the simulation. **c,e,g,i,** Individual frames from the simulations show the Nup98 FG condensate (grey) and the indicated probe (blue) with or without GR_10_ (orange). Interaction strengths of various simulation components were quantified using force field parameters: average interaction strength between a probe composed of 50% Y, F, or W and 50% I, V, L, A, or M with Nup98 FG domain, **j**; between the same probes and GR_10_, **k**; self-interactions for probes composed of 50% Y, F, or W and 50% I, V, L, A, or M (left), or 100% Y, F, or W (right), l. Data are shown as mean ± SD.

The impact of polyGR on probe partitioning is dependent on the chemistry of the amino acids. Probes composed of 50% aliphatic amino acids or 50% aromatic amino acids consistently show the largest impact from the addition of polyGR. Further, the identity of the aromatic amino acid dictates the overall behaviour of the system. Specifically, probes containing tyrosine showed the greatest increase in partitioning upon the addition of polyGR (Fig. 2b,c). In contrast, phenylalanine- and tryptophan-containing probes showed smaller changes from polyGR. This is likely due to the probes being already highly partitioned in the absence of polyGR, with phenylalanine-containing probes showing a slight increase in partitioning in the presence of polyGR, whereas tryptophan-containing probes show a slight decrease (Fig. 2d-g). Tryptophan-containing probes often form clusters with polyGR independently of Nup98 FG condensates, explaining their reduced partitioning. Despite the importance of aromatic amino acids in driving the ability of polyGR to alter probe partitioning, probes composed solely of aromatic amino acids showed relatively low partitioning into Nup98 FG condensates in the presence or absence of polyGR, again due to the formation of clusters outside of condensates (Fig. 2h,i).

To better explain these results, we used the parameters from the underlying force field to calculate the naive interaction strengths between different species (see *Methods*). For interaction strengths between probes with 50%:50% aliphatic:aromatic residues with Nup98, there was a direct correlation between each of their interaction strengths and the percentage of probe partitioned into the condensate in the absence of polyGR (Fig. 2j). Further, probes with tryptophan interacted more strongly with polyGR than those with tyrosine or phenylalanine (Fig. 2k), explaining the decrease in condensate partitioning of the tryptophan-containing probes with polyGR.

Finally, probes composed solely of aromatic residues showed substantially stronger self-interaction than those with both aromatic and aliphatic residues (Fig. 2l), highlighting how self-association can outcompete partitioning into Nup98 condensates.

Ultimately, our simulations suggest that a range of interaction strengths exist between i) a probe and itself, ii) a probe and Nup98, and iii) a probe and polyGR, and that the interplay among these interactions determines the ability of polyGR to impact partitioning of a given client across the nuclear pore. Furthermore, the strength of Nup98:Nup98 and Nup98:GR interactions can synergise or antagonise probe recruitment or exclusion. If the probe interacts too strongly with itself, self-assembly will outcompete putative Nup98 interactions, preventing partitioning. If the probe interacts strongly with polyGR, the client may form condensates with polyGR rather than partitioning into Nup98 condensates. Finally, if the probe interacts strongly with Nup98 at baseline, polyGR may be unable to further enhance its partitioning. Crucially, the multicomponent nature of this superficially simple system leads to a network of competing chemically specific interactions that governs how polyGR influences recruitment or exclusion into Nup98 condensates in a client-specific manner^70^.

### Continuum of condensate partitioning based on protein surface chemistry

In agreement with our simulations, previous work has demonstrated that the surface-exposed amino acids of client proteins determine their passive diffusion rates through nuclear pores^41^. Specifically, solvent-accessible hydrophobic residues on the protein surface promote partitioning into the FG phase and facilitate NPC passage. Clients with such residues can be referred to as FG-philic, while those lacking them as FG-phobic. Indeed, EGFP, whose FG partitioning is enhanced by polyGR, can be described as FG-philic as it contains several solvent-accessible hydrophobic residues enabling weak FG interactions^41^. Conversely, mCherry, which is unaffected by polyGR, predominantly contains FG-phobic residues on its surface - and thus can be considered FG-phobic^41, 56^.

To further test whether the impact of polyGR on partitioning into Nup98 FG condensates is dependent on client surface chemistry, we investigated the impact of polyGR on the partitioning of a GFP variant series developed by Frey *et al.*^41^. In this series, eight surface residues of the scaffold efGFP (enhanced folding of hydrophobic GFP) were mutated to modify interaction with FG domains to become more FG-philic or FG-phobic (Fig. 3a). Importantly, while these mutations alter the attraction to the FG phase based on surface chemistry, they do not significantly change its structure, molecular weight, or brightness (Extended Data Fig. 3a). We used this targeted efGFP surface mutant series to examine how surface amino acid chemistry shapes the effect of polyGR on client partitioning into our human Nup98 FG condensate system. We focused on solvent-exposed hydrophobic residues, which are the most important determinants of both nuclear pore passage rates and interaction with the FG phase^41^.

**Figure 3.**
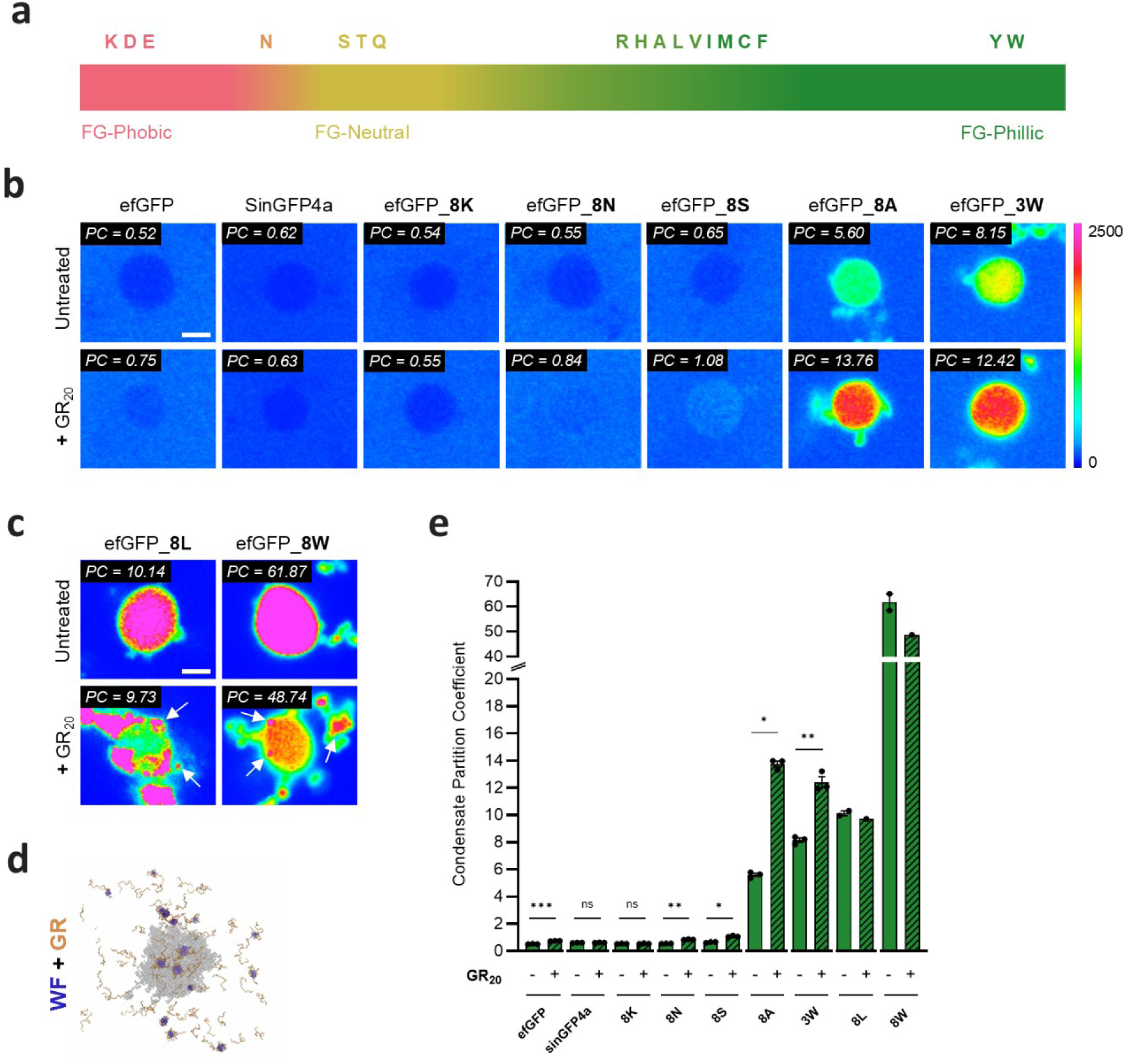
Client surface hydrophobicity determines polyGR-dependent modulation of FG phase partitioning. **a**, Schematic of the efGFP surface mutant series used to systematically vary client interactions with FG domains through defined changes in solvent-exposed hydrophobic and aromatic residues, without altering protein size or fold. **b, c, e,** Partitioning of efGFP and its surface chemistry variants into recombinant human Nup98 FG condensates in the absence or presence of GR_20_ - representative images and mean condensate partition coefficients (PC) (experiments performed separately but shown on same graph for ease of comparison). PolyGR modulates FG phase partitioning in a client-dependent manner: wild-type efGFP and the neutral or mildly FG-phobic variants efGFP_8S and efGFP_8N exhibit modest increases in partitioning upon GR_20_ exposure, whereas the FG-philic, hydrophobic variants efGFP_8A and efGFP_3W show strong enhancement. In contrast, the highly FG-phobic variants efGFP_8K and sinGFP4a are insensitive to GR_20_. Stats: two-tailed paired *t*-tests; n= 5 biological replicates, ≥5 condensates per replicate; ***p < 0.001; **p < 0.01; *p < 0.05; ns, not significant. For the extremely hydrophobic variants efGFP_8L and efGFP_8W, GR_20_ reduces FG-phase partitioning and promotes aggregation outside the FG condensate (arrows) (replicate numbers insufficient to perform statistical analysis). **d,** Coarse-grained simulation also demonstrates extra-Nup98 FG condensate accumulations of polyGR with probe; here showing the aromatic WF probe. Data are shown as mean ± SEM. Scale bars = 2 µm.

The partitioning of wild-type efGFP into Nup98 FG condensates was increased by polyGR (ΔPC = +0.23), as observed for EGFP. In examining the efGFP variant series, we found that the strength of the effect that polyGR has on partitioning was indeed dictated by the surface amino acid residues of each variant (Fig. 3b,c). The mildly FG-phobic mutant efGFP_8N and the neutral efGFP_8S variant showed a modest increase in their partitioning into condensates with polyGR (ΔPC = +0.29, ΔPC = +0.43, respectively). However, partitioning of the highly FG-phobic efGFP_8K variant was unchanged by polyGR (ΔPC = +0.01). We also tested the extremely FG-phobic variant sinGFP4a, which contains over 25 mutations that convert FG-philic residues to FG-phobic ones^41^. As with efGFP_8K, polyGR did not alter the partitioning of sinGFP4a into condensates (ΔPC = +0.01). In contrast, polyGR caused markedly increased partitioning of the FG-philic efGFP mutants efGFP_8A (ΔPC = +8.16) and efGFP_3W (ΔPC = +4.27). However, for the highly hydrophobic FG-philic variants efGFP_8L and efGFP_8W, polyGR led to a subtle (ΔPC = -0.41) or significant (ΔPC = -13.13) reduction, respectively, in partitioning, accompanied by aggregation of variants outside of condensates (Figure 3c,d). In agreement, we observed similar behaviour in our coarse-grained simulations where some of the aromatic amino acid-containing 10-mers that interact strongly with polyGR formed condensates outside of the Nup98 FG condensate (Figure 3d). This results in a biphasic response whereby polyGR increases the partitioning of increasingly FG-philic clients until a critical point after which the partitioning decreases, and extra-condensate accumulation and aggregation begins (Fig. 4c). Of note, the control *C9ORF72* DPR peptides polyGP and polyPA had no impact on efGFP or efGFP variant partitioning into Nup98 FG condensates (Extended Data Fig. 4b-e), and thus this is a specific effect of the arginine-rich polypeptide polyGR. These results suggest that polyGR causes a continuum of effects on client interaction with the nuclear pore transport barrier, depending on protein hydrophobicity and aromaticity.

**Figure 4.**
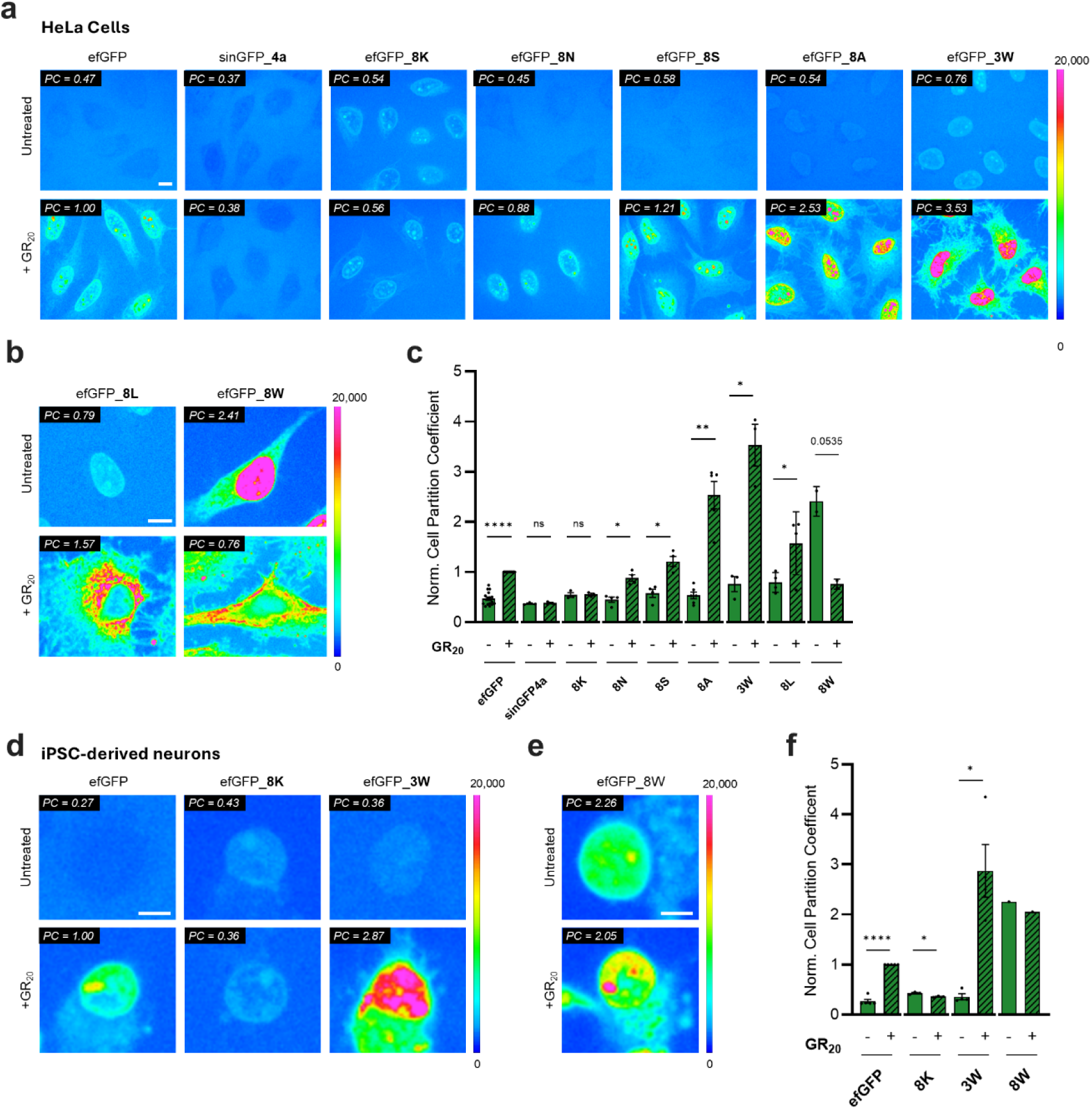
Condensate partitioning predicts polyGR-dependent disruption of passive nucleocytoplasmic transport in cell lines and human neurons. **a,b,c,** Representative endpoint heatmap images and cellular partition coefficients (PCs) of semi-permeabilised HeLa cells incubated with efGFP and variants for 4 min, either untreated or pre-treated with 10 µM GR_20_ for 30 min (experiments for each mutant were performed separately but assayed in parallel with efGFP, with and without GR_20_, within each biological replicate; PCs are normalised to efGFP+GR_20_ mean per replicate). PolyGR moderately increases nuclear import of wild-type efGFP, while having no effect on the highly FG-phobic variants sinGFP4a and efGFP_8K. Neutral variants efGFP_8N and efGFP_8S exhibit modest increases in nuclear import, whereas FG-philic hydrophobic variants, efGFP_8A and efGFP_3W, show pronounced enhancement. In contrast, highly hydrophobic FG-philic variants efGFP_8L and efGFP_8W undergo cytoplasmic aggregation upon polyGR treatment, with efGFP_8L displaying aggregation accompanied by a moderate increase in nuclear import, and efGFP_8W exhibiting aggregation together with a marked reduction in nuclear import. Stats: two-tailed paired *t*-tests for all except wild-type efGFP_0W, which was analysed using a one-sample *t*-test due to normalisation; n= 3 biological replicates for all efGFP variants except efGFP_8W (n=2); ****p < 0.0001, **p < 0.01, *p < 0.05; ns, not significant. Scale bars = 10 µm (a), 5 µm (b). **d, e, f,** Representative end-stage heatmap images after 4 min and cellular mean partition coefficients for efGFP variants in human iPSC-derived cortical neurons (experiments for each mutant were performed separately but assayed in parallel with efGFP, with and without GR_20_, within each biological replicate; PCs are normalised to efGFP+GR_20_ mean per replicate). Consistent with the HeLa cell data, polyGR increases nuclear import of wild-type efGFP, causes a modest decrease for the FG-phobic variant efGFP_8K, and produces a pronounced increase for the FG-philic hydrophobic variant efGFP_3W. Stats: two-tailed paired *t*-tests for all except wild-type efGFP_0W, which was analysed using a one-sample *t*-test due to normalisation; n= 3 biological replicates; ****p < 0.0001, *p < 0.05. For the extremely hydrophobic variant GFP_8W, GR_20_ results in a mild reduction in nuclear import accompanied by cytoplasmic aggregation (replicate numbers insufficient to perform statistical analysis). Scale bar = 10 µm for HeLa cells; 5 µm for human iPSC-derived cortical neurons.

### Condensate partitioning predicts passive nucleocytoplasmic transport disruption by polyGR in human cell lines and iPSC-derived neurons

Motivated by our *in silico* and *in vitro* results, we next sought to test whether client surface attributes predict the impact of polyGR on protein passive diffusion through intact nuclear pores. We again used the efGFP surface mutant series in combination with the classic permeabilised cell nuclear import assay in the human HeLa cell line (Fig. 4a-c). Wild type efGFP showed a moderate increase in nuclear import following polyGR treatment (ΔPC = +0.53). As in our *in vitro* condensate assay, the FG-phobic mutants efGFP_8K (ΔPC = 0.01) and sinGFP4a (ΔPC = +0.01) did not show any significant increases in nuclear influx following polyGR treatment. The slightly FG-phobic mutant efGFP_8N and the FG-neutral efGFP_8S variant showed mild polyGR-induced increase in import (ΔPC = +0.43 and +0.64, respectively). As expected, polyGR treatment resulted in a marked enhancement in nuclear import of the hydrophobic efGFP mutants efGFP_8A (ΔPC = +2.00) and efGFP_3W (ΔPC = +2.77). The highly hydrophobic and FG-philic efGFP variants efGFP_8L and efGFP_8W showed robust nuclear import in untreated cells. However, polyGR treatment no longer dramatically enhanced import, but led to a smaller increase in efGFP_8L partitioning (ΔPC = +0.78) or a decrease in efGFP_8W (ΔPC = -1.65). Notably, addition of polyGR causes the cytoplasmic aggregation of both 8L and 8W efGFP variants (Fig. 4b). The results of our cell assays mirror the findings in our *in silico* and *in vitro* condensate assays with a biphasic effect of polyGR on proteins with increasing surface hydrophobicity, increasing and then decreasing nucleocytoplasmic transport, and at the higher end resulting in cytoplasmic aggregation. Thus, our data provide strong evidence that protein surface hydrophobicity predicts the vulnerability and fate of cellular proteins to disease-associated polyGR polypeptides.

As the *C9ORF72* mutation and its associated aberrant peptides are causative of neurodegenerative diseases, we sought to corroborate this working model in human neuronal cells. Here, we used a control reference induced pluripotent cell line (KOLF2.1J^71^) differentiated using the transcription factor neurogenin-2 into predominantly excitatory cortical-like glutamatergic neurons (Extended Data Figure 5a), which were aged for 14 days and subjected to our permeabilised cell nuclear import assay.

Consistent with the HeLa cell line, polyGR caused a marked increase in the nuclear import of wild-type efGFP in neurons (ΔPC = +0.77) and no effect on mCherry (ΔPC = 0.00; Extended Data Figure 5b,c). Likewise, using select mutants from the efGFP series, polyGR exerted no impact on the FG-phobic efGFP_8K (ΔPC = -0.07), a significant enhancement of the FG-philic and hydrophobic efGFP_3W (ΔPC = +2.51) compared to wild-type efGFP (ΔPC = +0.73), and cytoplasmic accumulation and reduction in partitioning of the highly FG-philic efGFP_8W (ΔPC = -0.21) (Figure 4d-f), corroborating that a biphasic effect of polyGR on transport is conserved in human neurons.

These results demonstrate that the *C9ORF72* polypeptide polyGR preferentially enhances the nuclear import of proteins with hydrophobic surface residues that have an affinity for the nuclear pore FG phase, while having little or no effect on FG-phobic proteins. Importantly, our results demonstrate that solvent-exposed amino acids are the predominant factor determining the effect of aberrant peptides on passive nucleocytoplasmic transport rates. However, a continuum exists in polyGR behaviour from promoting nuclear transport of moderately FG-philic proteins to driving cytoplasmic aggregation of highly FG-philic proteins. We validate that these findings are consistent between immortalised human cell lines and iPSC-derived neurons. Finally, we establish that cellular nucleocytoplasmic transport modulation by aberrant peptides can be predicted using *in silico* and *in vitro* condensate modelling of the nuclear pore FG phase.

### PolyGR binds the nuclear pore central channel in human cells

A key assumption of our surface property model is that polyGR peptides directly interact with NPCs in cells, particularly the FG-rich permeability barrier. Previous work in isolated giant nuclei from *Xenopus laevis* oocytes showed that a fluorescently labelled polyPR (the other arginine-rich DPR) bound to the NPC central channel^72^. However, it is unknown whether the same holds for polyGR in intact human nuclear pores. To determine whether polyGR directly interacts with the NPC in human cells, we turned to super-resolution microscopy.

Using two-colour STORM, we simultaneously visualised polyGR polypeptides (labelled with the dye ATTO655) and the NPC scaffold component NUP96 (endogenously tagged with mEGFP, detected with anti-GFP nanobodies by DNA-PAINT) in U2OS cells (Fig. 5a). Importantly, ATTO655 dye alone did not localise to nuclear pores (Fig. 5b; Extended Data Fig. 7a), although it could penetrate cells (Extended Data Fig. 7c,d). In contrast, ATTO655-polyGR exhibited distinct localisation in close proximity to NPCs, not co-localising with the structural component Nup96, but instead in surrounding regions, including within the central channel (Fig. 5b; Extended Data Fig. 7b), strongly suggesting that polyGR can access and bind to FG-containing NPC domains. Taken together, our results support the disruption of passive nucleocytoplasmic transport by polyGR through targeting and binding of intrinsically disordered NPC components in human cells.

**Figure 5.**
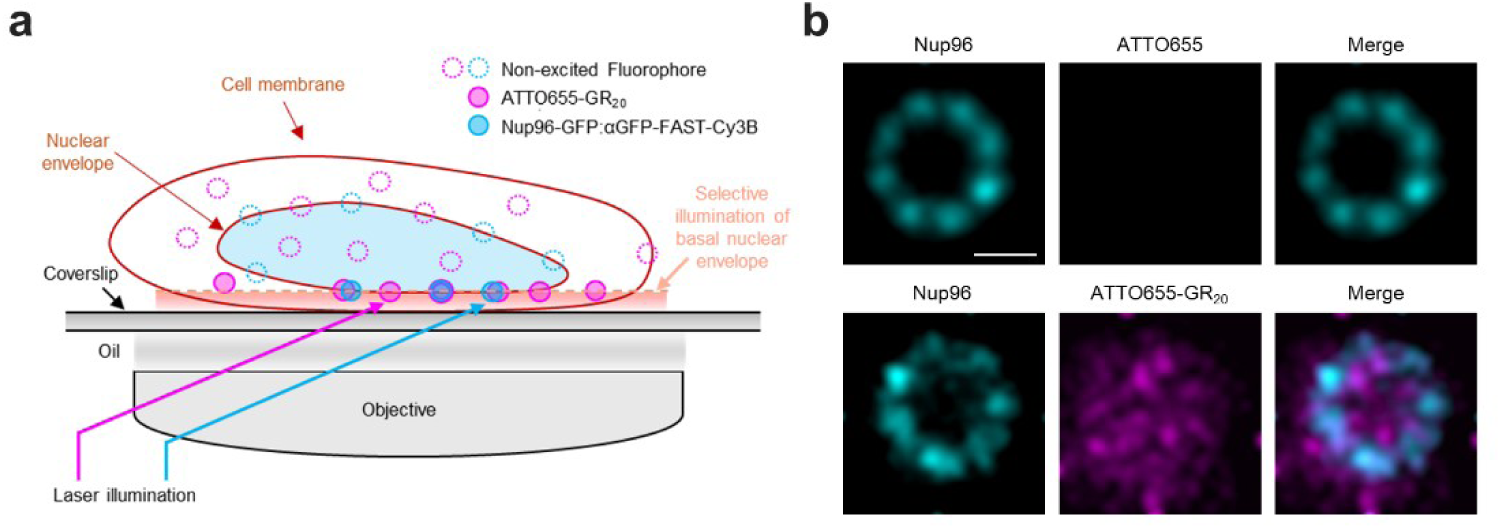
PolyGR directly associates with the nuclear pore complex in human cells. **a,** Schematic of the experimental design for assessing polyGR association with nuclear pores using two-colour STORM microscopy in U2OS-CRISPR–NUP96–mEGFP cells. Angled beam imaging was used to selectively excite single molecules of ATTO655-labelled GR_20_ or ATTO655 dye alone and Nup96-mEGFP labelled with anti-GFP nanobodies by DNA-PAINT at the basal nuclear envelope. **b,** Averaged NPC localisations from STORM imaging of fixed cells showing that ATTO655 dye alone does not localize to NPCs (upper panel), whereas ATTO655–GR_20_ exhibits localisation in close proximity to NPCs, not co-localising with the structural component Nup96, but instead in surrounding regions including within the central channel. Scale bars = 50 nm.

## Discussion

In this study we demonstrate that polyGR - an aberrant neurotoxic polypeptide derived from the *C9ORF72* repeat expansion mutation in ALS/FTD – pathologically remodels the physiochemical selectivity of the nuclear pore. By directly engaging the FG phase, polyGR alters passive nucleocytoplasmic transport in a manner dictated predominantly by client surface chemistry rather than size. This produces biphasic transport behaviour in which FG-phobic proteins are unaffected, but progressively more FG-philic and hydrophobic proteins initially exhibit enhanced nuclear pore passage, until a critical threshold is reached, beyond which highly hydrophobic proteins instead experience transport suppression, cytoplasmic accumulation, and aggregation (Fig. 6).

**Figure 6.**
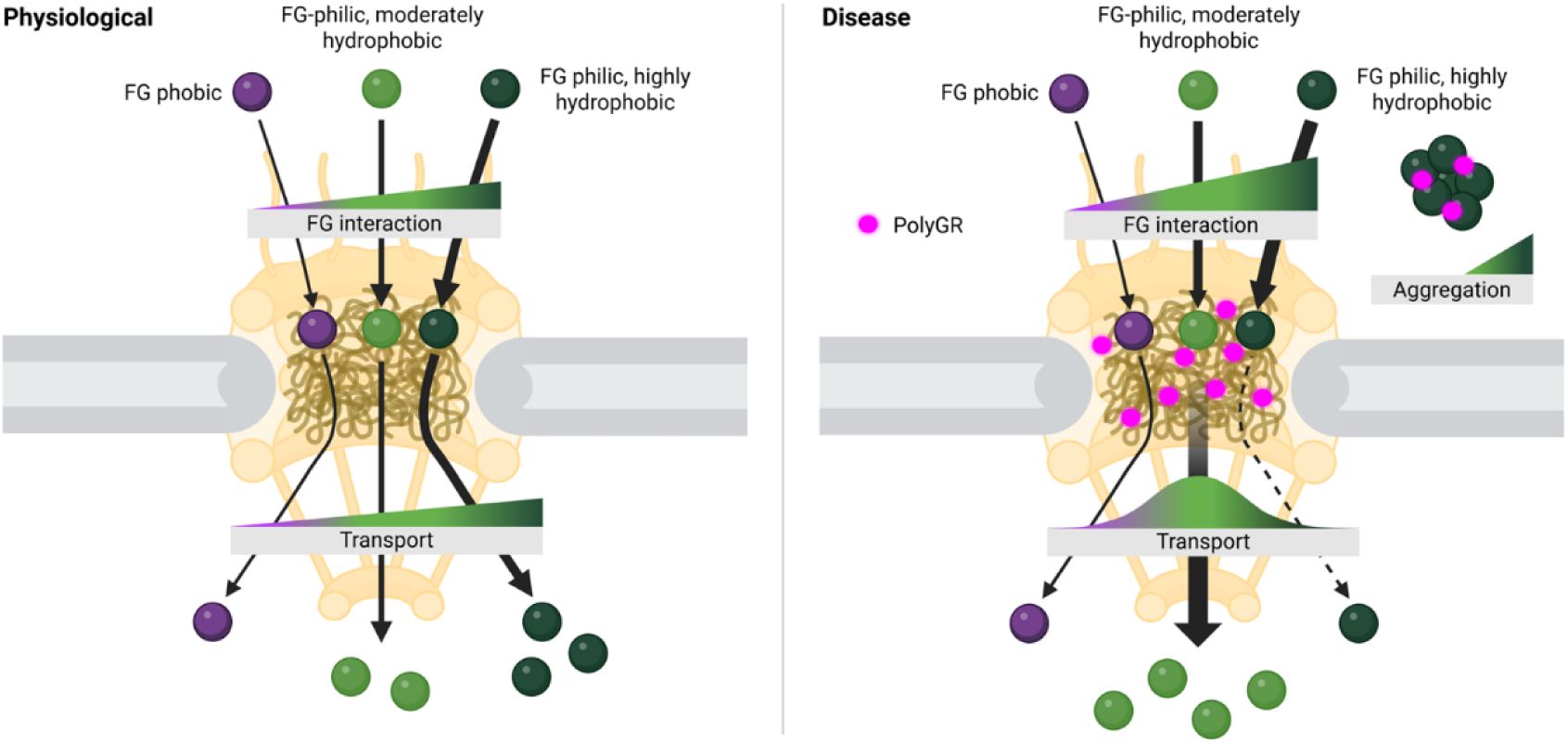
PolyGR retunes physicochemical rules governing passive nuclear pore transport. Under basal conditions, passive transport through the nuclear pore complex (NPC) is governed by interactions between transiting clients and the FG-rich permeability barrier. Clients with low affinity for FG domains diffuse slowly or are excluded, whereas FG-philic clients partition into and traverse the pore efficiently, with the strength of these interactions determined by surface hydrophobicity. In the presence of the arginine-rich polypeptide polyGR, the physicochemical environment of the FG barrier is altered. PolyGR dynamically retunes NPC selectivity in a cargo-dependent manner: FG-phobic proteins remain largely unaffected, moderately FG-philic proteins exhibit enhanced nuclear passage, and highly hydrophobic FG-philic proteins are diverted from the pore, leading to reduced nuclear import and cytoplasmic accumulation or aggregation. Rather than uniformly blocking or enhancing transport, polyGR reshapes nuclear pore selectivity, generating a continuum of outcomes determined by cargo surface chemistry.

Under physiological conditions, the FG-rich nucleoporin Nup98 plays a crucial role in mediating selective nucleocytoplasmic transport^38^, with its disordered FG domain forming a chemically tuned permeability barrier^36–38, 41, 50^. Consistent with established surface-chemistry principles governing FG-phase interactions^41^, the client partitioning patterns we observe for isolated Nup98, both *in silico* and *in vitro*, are preserved within intact nuclear pores in cells. Importantly, using super-resolution imaging, we demonstrate that polyGR binds within the FG-filled central channel of intact nuclear pores in cells, confirming that these interactions occur within the native NPC environment. In parallel, we show that the aberrant polypeptide polyGR directly engages FG-rich environments in vitro, including recombinant Nup98 FG condensates. Systematic substitution of FG motifs reveals that aromatic residues within the human Nup98 FG domain, particularly phenylalanine, are the primary mediators of polyGR binding: F>Y substitutions preserve strong interactions, whereas F>L or F>S progressively weaken them. These data support a multivalent interaction network dominated by aromatic and hydrophobic contacts, likely involving arginine guanidinium-pi interactions, with additional contributions from electrostatic and hydrogen bonding interactions. Collectively, these findings establish that polyGR exploits the same chemical features that normally govern FG phase interactions and transport of clients through the nuclear pore.

Previous work has shown that polyGR can accelerate passive nuclear import in a size-dependent manner^26^. We confirm this effect; however, our data demonstrate that size alone is insufficient to predict the impact of polyGR on a given protein. Instead, polyGR exerts selective, surface-chemistry–dependent effects on clients of comparable molecular weight. For example, whereas the transport of the efGFP_8K and SinGFP4a variants is completely unaffected by polyGR, the 3W and 8A variants exhibit markedly enhanced transport, despite all four proteins having similar molecular weights. These observations demonstrate that client surface properties are key determinants of susceptibility to polyGR-mediated modulation of nucleocytoplasmic transport. The FG phase of the nuclear pore functions as a selective solvent^37, 39, 73^, in which client surface amino acids, particularly aromatic and hydrophobic residues that interact strongly with the FG repeat network, govern partitioning and transport^39, 41, 74^. Our data show that this interaction-dependent transport behaviour is pathologically amplified by polyGR, producing a biphasic response in which progressively FG-philic proteins initially exhibit enhanced nuclear pore passage, until a threshold is reached beyond which transport is suppressed and clients are diverted toward cytoplasmic accumulation and aggregation. In doing so, polyGR does not simply shift the existing client–NPC transport continuum but introduces an additional aggregation-prone regime.

PolyGR-mediated perturbation of FG Nup interactions is likely to influence nucleocytoplasmic transport through multiple, non-mutually exclusive mechanisms. Fluorescence recovery after photobleaching reveals no detectable change in the mobility of nuclear transport receptors within Nup98 FG condensates, arguing against a global “loosening” of the FG meshwork. Instead, condensate assays and simulations indicate that polyGR can directly compete with client proteins for FG-motif interactions whilst simultaneously engaging solvent-exposed residues on client surfaces, thereby increasing their local concentration within the FG phase. In this context, FG motifs occupied by polyGR may act as transient recruitment sites for chemically compatible clients, tuning interaction lifetimes during transit. For weakly FG-philic clients, polyGR may introduce additional contacts, such as electrostatic interactions, which enhance recruitment and favourably alter interaction dynamics. Conversely, by partially occupying FG binding sites, polyGR may reduce the effective dwell time of certain FG-client interactions, thereby accelerating client diffusion through the nuclear pore. This dual role as both recruiter and interaction modulator provides a mechanistic explanation for the increased accumulation of FG-philic relative to FG-phobic clients within the FG phase observed *in vitro* and *in silico*, as well as the enhanced passive transport measured in cells. Accordingly, we propose that polyGR acts as a context-dependent “molecular lubricant” within the FG phase, facilitating the transit of chemically compatible clients and tuning barrier selectivity. This stands in contrast to a previously proposed model whereby the arginine DPRs were suggested to bind and occlude the pore^72^. Consistent with our model, the polyGR-induced increase in passive nuclear import (here, of the inert 10 kDa dextran) is unaffected by co-treatment with the highly FG-philic efGFP_3W variant, demonstrating that polyGR does not block transport, even under conditions of strong FG engagement (Extended Data Fig. 5d,e).

In contrast to the enhancement state described above, the transition from enhanced to suppressed nucleocytoplasmic transport, together with aggregation onset, likely arises via distinct mechanisms for the two highly hydrophobic client variants examined. For the aromatic-rich efGFP_8W variant, reduced partitioning in the presence of polyGR is consistent with co-assembly in the dilute phase, supported by simulations revealing strong cation–pi interactions between polyGR and aromatic residues that could stabilize polyGR-efGFP_8W complexes. Such interactions could sequester efGFP_8W, thereby reducing the pool of freely available protein competent for nuclear pore passage. In this respect, the effect of polyGR on efGFP_8W mirrors previously reported interactions between arginine-rich DPRs (including polyGR) and nuclear transport receptors, which are among the most hydrophobic soluble proteins^41, 73^. Here, DPR binding promotes phase separation and insolubility of transport receptors, rendering them unavailable for and thereby disrupting active (receptor-mediated) nucleocytoplasmic transport^25, 26, 75^; although the effect on active transport has not been uniform across studies^76^. In contrast, the aliphatic-hydrophobic efGFP_8L variant does not contain an aromatic structure and thus lacks such strong cation–pi interactions with polyGR, suggesting a distinct mechanism. Here, polyGR likely alters the effective chemistry of the FG phase, shifting it toward a more arginine-rich and less hydrophobic environment that disfavours entry of strongly aliphatic sequences. Reduced FG-phase partitioning of efGFP_8L would increase its cytoplasmic concentration and promote concentration-driven aggregation, an effect further exacerbated by its intrinsic propensity for self-association. Together, these divergent responses of aromatic versus aliphatic clients underscore limitations of current coarse-grained simulation frameworks, including those used here, which tend to underestimate the impact of hydrophobicity and thus fail to fully capture aliphatic-driven self-assembly, and underscore the necessity of cross-validating computational findings in biological systems. Also, it is likely that there is a critical point of FG affinity beyond which transport is no longer favourable even in physiological conditions, however this limit was not reached in our study. Together, our findings indicate that polyGR reshapes passive nucleocytoplasmic transport in a chemically encoded, client-specific manner, naturally raising the question of how endogenous proteins with complex surface chemistries behave within polyGR-perturbed nuclear pores.

In relation to disease, these mechanistic distinctions focus attention on a central unresolved question - why certain proteins are selectively vulnerable to mislocalisation in ALS/FTD. Our finding that polyGR modulates passive nucleocytoplasmic transport in a client chemistry-dependent manner provides a mechanistic framework to explain this selectivity. This is particularly relevant to the cytoplasmic mislocalisation of TDP-43 in *C9ORF72*-ALS/FTD. Although TDP-43 is actively imported into the nucleus, it is passively exported to the cytoplasm^77, 78^, rendering its steady-state localisation sensitive to interactions within the FG phase of the nuclear pore. Notably, TDP-43 is enriched in hydrophobic residues across its C-terminal intrinsically disordered region, N-terminal domain, and RNA recognition motifs^78^. Because transport of intrinsically disordered proteins through the NPC depends strongly on the balance of hydrophobic and charged residues rather than size alone^79, 80^, these features are likely to influence TDP-43 diffusion through the FG phase and render it susceptible to polyGR–mediated perturbation. This may suggest that polyGR-induced reprogramming of FG-phase chemistry exacerbates passive nuclear export of TDP-43, promoting its cytoplasmic accumulation and aggregation – an important hypothesis to test in future studies. Together with prior reports of polyGR-induced sequestration of nuclear transports^25, 26^ and *C9ORF72*-associated increases in nuclear pore component turnover^81, 82^, our findings point to multiple, convergent routes to nuclear pore dysfunction in *C9ORF72*-ALS/FTD that ultimately lead to TDP-43 pathology.

More broadly, our work demonstrates how disease-associated aberrant polypeptides disrupt not only the material properties of biomolecular condensates but also their emergent functions, with the nuclear pore providing a prominent example. Through an integrated combination of *in silico*, *in vitro* and cellular approaches, we show that the *C9ORF72*-derived polypeptide polyGR acutely reprograms client partitioning into the FG phase of the nuclear pore in a surface chemistry-dependent and biphasic manner, with hydrophobicity emerging as a dominant determinant. These shifts in FG phase partitioning quantitatively predict altered nucleocytoplasmic transport and downstream cytoplasmic aggregation. Consistent with the broader relevance of surface residue-FG interactions in human disease, similar principles have recently been implicated in HIV-mediated nuclear import^39, 56, 57^. Together, this work establishes a general framework in which multi-component condensate interaction mapping can be used to link altered condensate chemistry to functional impairment, providing mechanistic insight into cellular dysregulation in disease. Applied here to the nuclear pore, this framework reveals previously unappreciated contributors to disrupted nucleocytoplasmic homeostasis that drive neurodegeneration in *C9ORF72*-ALS/FTD.

## Methods

### Plasmids

The wild-type human Nup98 FG domain and the F>Y, F>L and F>S variant plasmids were synthesised by GeneArt (Thermo Fischer Scientific). The efGFP mutant series^41^ and proteases^83^ required for their purification were kindly provided by Dirk Gorlich (Max Planck Institute for Multidisciplinary Sciences). All other plasmids were acquired from AddGene. Plasmids used for recombinant protein purification are listed in Supplementary Table 1.

### Recombinant protein expression and purification

Recombinantly purified SinGFP4a^41^ and Alexa488 tagged NTF2^50^ proteins were a kind gift from Dirk Gorlich (Max Planck Institute for Multidisciplinary Sciences). Recombinant EGFP was obtained from Cambridge Bioscience (STA-201) and mCherry from Abcam (ab199750).

### Human Nup98 FG domain and F>Y, F>L, and F>S variants

The expression and purification of wild-type Nup98 FG domain (residues 1–499) and the F>Y, F>L, and F>S variants was adapted from Frey *et al.*^41^. Hisₓ₁₄-tagged human Nup98 FG domain (residues 1–499) and Hisₓ₁₄-tagged F>Y, F>L, and F>S variants were expressed in BL21 Star™ (DE3)pLysS One Shot™ Chemically Competent E. coli (Thermo Fisher Scientific, C602003). Starter cultures were grown in Luria Broth (LB; VWR International, 85856.5000) containing ampicillin (100 μg/mL, Sigma-Aldrich, A5354) and chloramphenicol (34 μg/mL, BioServ, BS-0258A-5G) overnight at 37 °C, then diluted 1:100 into Terrific Broth (TB; Merck Millipore, 1.01629.0500) supplemented with 0.4% (v/v) glycerol (Thermo Fischer Scientific, 15514011), and grown at 37 °C to OD₆₀₀ ≈ 1. Expression was induced with 400 μM IPTG (Thermo Fischer Scientific, AM9464) for 4 h at 30 °C. Cells were harvested by centrifugation (5000 rpm, k = 8234, 4 °C) and stored at –80 °C. The human Nup98 FG domain and F>Y variant pellets were thawed on ice and resuspended in 50 mM Tris-HCl pH 7.5 (Sigma Aldrich, 252859), 300 mM NaCl (Thermo Fischer Scientific, A57006), 1 mg/mL lysozyme (Sigma Aldrich, L6876), 1 mM PMSF (Thermo Fisher Scientific, 36978), 0.1 μg/mL DNase I (Roche, 10104159001), and EDTA-free protease inhibitors (Sigma Aldrich, 06538282001). Cells were lysed by mild sonication. Lysates were centrifuged (9920 rpm, k = 2600, 15 min, 4 °C), and the resulting pellet was washed and resuspended in 50 mM Tris-HCl pH 7.5, 300 mM NaCl, 5 mM DTT (Thermo Fischer Scientific, 15508013), 1 mM PMSF, and EDTA-free protease inhibitors. After a second centrifugation under the same conditions, the supernatant was discarded, and FG domains were extracted from the pellet by resuspension in 6 M guanidine-HCl (Sigma Aldrich, G3272), 50 mM Tris-HCl pH 8.0, 1 mM imidazole pH 7.5, 10 mM DTT, 1 mM PMSF, and EDTA-free protease inhibitors. The extracted material was clarified by ultracentrifugation (38,000 rpm, k = 135, 4 °C). F>L and F>S variants were thawed and lysed directly in 6 M guanidine-HCl, 50 mM Tris-HCl pH 8.0, 1 mM imidazole pH 7.5, 10 mM DTT, 1 mg/mL lysozyme, 1 mM PMSF, 0.1 μg/mL DNase I, and EDTA-free protease inhibitors, followed by sonication and ultracentrifugation as above. The Nup98 FG domain and F>Y, F>L, and F>S variant supernatants were applied to a Ni-Agarose column (Qiagen, 30210) for 2 h at 4 °C. The column was sequentially washed in 6 M GuHCl, 50 mM Tris-HCl pH 8.0, 10 mM DTT, with imidazole pH 7.5 added in a gradient from 1–40 mM. Proteins were eluted from the column with 4 M GuHCl, 50 mM Tris-HCl pH 8.0, 500 mM imidazole pH 7.5 (bioworld, 40120857), and 10 mM DTT. Eluates were buffer-exchanged into 6 M GuHCl, 50 mM Tris-HCl pH 8.0, 10 mM DTT using PD-10 desalting columns (Cytiva, 17085101) and further purified by size-exclusion chromatography under denaturing conditions using either a HiLoad 26/600 Superdex 200 pg column (human Nup98 FG domain) or a HiLoad 16/600 Superdex 200 pg column (F>Y, F>L, and F>S variants), both equilibrated and run in 6 M GuHCl, 50 mM Tris-HCl (pH 8.0), and 10 mM DTT. Protein-containing fractions were pooled, concentrated, exchanged into ddH₂O + 0.08% TFA (VWR International, 0993), and lyophilised. As the absorbance of FG domains at 280nM is extremely low^84^ and GuHCl is not compatible with colorimetric protein concentetion assays the most accurate way to determine FG Nup and FG Nup mutant concentration is via gravimetry. Lyophilised protein was weighed and then dissolved to 400 μM in 2 M GuHCl, 100 mM Tris-HCl pH 8.0, aliquoted, flash-frozen in liquid nitrogen, and stored at –80 °C.

### efGFP mutant series

Expression and purification of the efGFP mutant series was adapted from Frey *et al.*^41^. Plasmids were transformed into NEB Express (MiniF LacIq) *E. coli* (NEB, C3037I). Overnight LB cultures grown from single colonies at 37 °C were diluted 1:100 into Terrific Broth containing 0.4% (v/v) glycerol and incubated at 37 °C until OD₆₀₀ ≈ 2. Protein expression was induced with 0.2 mM IPTG at 18 °C overnight. Cells were harvested by centrifugation (5000 rpm, k = 8234, 4 °C) and pellets stored at –80 °C. For extraction, pellets were thawed on ice and lysed in 50 mM Tris-HCl pH 7.5, 300 mM NaCl, 20 mM imidazole pH 7.5, 5 mM DTT, and protease inhibitors (EDTA-free tablet), followed by sonication. Lysates were clarified by ultracentrifugation (38,000 rpm, k = 135, 4 °C), and the supernatant was applied to a Ni-Agarose column for 2 h at 4 °C. The column was then washed sequentially in the following buffers – (i) 50mM Tris pH8; 300mM NaCl; 25mM Imidazole; 10mM DTT; (ii) 50mM Tris pH8, 1000mM NaCl, 5mM DTT; (iii) 50mM Tris pH8, 100mM KCl (Thermo Fischer Scientific, AM9640G), 5mM MgCl2 (Thermo Fischer Scientific, AM9530G), 1mM ATP (Thermo Fischer Scientific, R0441), 5mM DTT; and finally (iv) 50 mM Tris pH8, 300 mM NaCl, 250 mM sucrose (Sigma Aldrich, 84097), 2 mM DTT. On-column cleavage^83, 85^ was performed by applying protease buffer (50 mM Tris-HCl pH 8.0, 300 mM NaCl, 250 mM sucrose, 2 mM DTT) containing the appropriate protease (bdSENP1 or scSUMOstar) in a volume equal to the resin bed. The buffer was allowed to enter slowly, leaving a small volume above the resin to maintain hydration, and the column incubated at 4 °C overnight. Cleaved efGFP variants were collected as the primary flow-through. Remaining protein was recovered by washing the column with incremental additions of protease buffer without protease. Fractions exhibiting the most intense fluorescence were pooled. The efGFP proteins were further purified on an ÄKTA pure system using a HiLoad 16/600 Superdex 75/200 pg column depending on the predicted molecular weight of the efGFP variant^41^Protein-containing fractions were pooled, and protein concentration was determined by measuring absorbance at 488 nm (GFP absorption maximum) on a NanoDrop™/OneC, and molar concentration was calculated using the Lambert-Beer law:

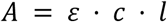

Where *A* is absorbance, *C* is the molar concentration of a GFP monomer unit, *d* is path length, ε is the extinction coefficient, which is 55000 M^-1^cm^-1^ (of a GFP monomer unit)

To determine the concentration, the equation was rearranged as follows:

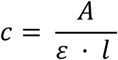

### bdSENP1, scSUMOstar and superTEV proteases

SUMO-specific proteases from Brachypodium distachyon (bdSENP1) and Saccharomyces cerevisiae (scSUMOstar) were purified following Frey *et al.*^83^. Plasmids encoding His₁₄-TEV-bdSENP1 and His₁₄-TEV-scSUMOstar were transformed into NEB Express (MiniF LacIq) E. coli. Single colonies were grown overnight in LB at 37 °C and diluted 1:100 into Terrific Broth supplemented with 0.4% (v/v) glycerol. Cultures were grown at 37 °C to OD₆₀₀ ≈ 2.0, induced with 0.2 mM IPTG, and incubated overnight at 18 °C. Cultures were supplemented with 5 mM EDTA prior to harvesting by centrifugation (5000 rpm, k = 8234, 4 °C), and cell pellets were stored at –80 °C. For extraction, pellets were thawed on ice and resuspended in lysis buffer (290 mM NaCl, 45 mM Tris-HCl pH 7.5, 4.5 mM MgCl₂, 10 mM DTT, 15 mM imidazole) and lysed by sonication. Lysates were clarified by ultracentrifugation (200,000×g, 1 h, 4 °C). Supernatants were applied to a Ni-Agarose column for 1 h at 4 °C. The bdSENP1/scSUMOstar proteases were eluted with 290 mM NaCl, 45 mM Tris-HCl pH 7.5, 4.5 mM MgCl₂, 10 mM DTT, 400 mM imidazole. His₁₄-TEV tags were removed by incubating the eluate overnight at room temperature with His-tagged Super TEV protease at a 1:30 molar ratio (protease:substrate). The mixture was then applied to a HiLoad 26/600 Superdex 200 pg column on an ÄKTA pure system, pre-equilibrated in 290 mM NaCl, 45 mM Tris-HCl pH 7.5, 4.5 mM MgCl₂, 5 mM DTT. To remove residual TEV protease, pooled peak fractions were passed twice over Ni-Agarose columns at 4 °C, with 1 h incubation between each pass. Protein concentration of the final flow-through fractions was determined using the Bradford assay (Bio-Rad, 5000205). Purified proteases were snap-frozen in liquid nitrogen and stored at –80 °C.

Super TEV (for cleaving bdSENP1 and scSUMOstar proteases) was purified as previously described^86^ with minor modifications. E. coli BL21(DE3) CodonPlus-RIL cells (Agilent, 230245) harbouring pDZ2087 (Addgene, #92414) were grown overnight in LB medium supplemented with ampicillin (100 μg/mL) and chloramphenicol (30 μg/mL) at 37 °C with shaking. Cultures were diluted 1:40 into fresh LB medium containing antibiotics and 0.2% (w/v) glucose (Sigma Aldrich, G7528) and incubated at 37 °C until mid-log phase (OD₆₀₀ ≈ 0.5). Protein expression was induced with 1 mM IPTG at 30 °C for 4–5 h. Cells were harvested by centrifugation (5000 rpm, 10 min, 4 °C, k-factor: 8234), and pellets were stored at −80 °C until further use. Frozen cell pellets were resuspended in lysis buffer (50 mM sodium phosphate pH 8.0, 200 mM NaCl, 25 mM imidazole) and lysed by sonication. Cell debris and nucleic acids were removed by 0.1 % polyethylenimine treatment (Sigma Aldrich, P3143) and centrifugation (15,000 × g, 30 min). The clarified lysate was applied to a Ni-NTA column, washed extensively, and eluted with an imidazole gradient. Fractions containing TEV protease were pooled, supplemented with 2 mM EDTA (Thermo Fischer Scientific, 15575020) and 5 mM DTT, and concentrated. The sample was further purified by size-exclusion chromatography on a HiLoad 16/600 Superdex 200 pg column. Pooled peak fractions were concentrated, filtered, aliquoted, flash-frozen in liquid nitrogen, and stored at −80 °C. The protein concentration was determined via Bradford assay (BioRad, 5000205), according to the standard protocol.

### Nup98 FG domain and mutants gel electrophoresis

Purity of the purified human Nup98 FG domain and F>Y, F>L, and F>S variants was assessed following Palmer & Wingfield^87^. To remove guanidine-HCl, samples were precipitated with isopropanol: 25 μL of protein was mixed with 225 μL cold 100% isopropanol and incubated at –80 °C for 10 min. Samples were centrifuged in a bench-top centrifuge at 15,000×g for 5 min at 4 °C, and the supernatant discarded. Pellets were resuspended in 250 μL cold 90% isopropanol (Thermo Fischer Scientific, BP2618-500) and centrifuged again under the same conditions. The final pellet was resuspended in 25 μL 1x SDS sample buffer for subsequent analysis. Protein purity was confirmed via gel electrophoresis using NuPAGE™ Bis-Tris Mini Protein Gels, 12%, 1.0 mm (Thermo Fischer Scientific, NP0342BOX) followed by Coomassie staining (abcam, ab119211).

### DPR peptide synthesis

Unlabelled 40mer DPR peptides for polyGP (GP_20_), polyPA (PA_20_), polyGR (GR_20_), Alexa Fluor 488 polyGR (AF488-GR_20_), Alexa Flour 488 polyPA (AF488-PA_20_) and ATTO 655 labelled polyGR (ATTO655-GR_20_) were synthesized by ThermoFisher Scientific. Lyophilised peptides were resuspended in HyClone™ HyPure molecular biology grade water (Cytiva, SH30538.FS) according to manufacturer’s instructions at a 1 mM concentration, then snap frozen in liquid nitrogen and stored at -80°C.

### *In vitro* FG phase permeation assays

The Nup98 FG domain stock solution (400 µM in 2 M GuHCl) was diluted rapidly to a final concentration of 5 µM in either assay buffer (TBS; 50 mM Tris/HCl pH 7.5, 150 mM NaCl) only or assay buffer + 100 µM DPR peptide (from a 1mM DPR stock). This rapid dilution of the GuHCl to negligible levels causes the almost instantaneous phase separation of the Nup98 FG domain into condensates. The Nup98 FG domain containing suspensions were then mixed thoroughly with fluorescent substrate. All efGFP mutants and EGFP were used at 1 μM, mCherry at 3 μM. Prior to imaging, the Nup98 FG condensates were left to sediment to the bottom of the slide (ibiTreat surface-modified µ-Slide 15-well 81506; ibidi) for 30 minutes. For all partitioning experiments (EGFP, mCherry, efGFP variants), unlabelled DPRs were used.

### Confocal laser-scanning microscopy of FG condensates

Images of condensates and their interaction with EGFP, efGFP variants, NTF2-Alexa488 and mCherry were acquired using Nikon AX/AX R inverted confocal microscopes with a 63x oil objective with a 488 or 561 nm laser at room temperature. All efGFP variants and EGFP were scanned on the same confocal settings (laser power and offset), except for efGFP_8L and efGFP_8W which were scanned at lower settings due to detector saturation but kept the same across technical and biological replicates.

### Quantification of partition coefficients

Partition coefficients (PCs) for fluorescent clients within each condensate were determined using the following relationship:

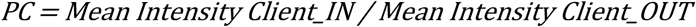

ΔPC represents the difference in partitioning coefficient between the presence and absence of polyGR:

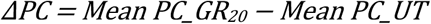

### Fluorescence recovery after photobleaching (FRAP)

NUP98 FG domain condensates were first formed either in the presence or absence of 100 µM GR_20_ peptide, followed by incubation with 1 µM NTF2-Alexa488. Samples were photobleached using a 488 nm laser with a 100 ms bleaching pulse at room temperature. Photobleaching was targeted to circular regions ∼1 µm in diameter within the FG phase. However, due to laser blurring and diffusion during bleaching, the actual bleached regions were approximately ∼2 µm in diameter. Following photobleaching, fluorescence recovery of Alexa488 was imaged every 10 seconds over a period of 10 minutes. Raw data were corrected for acquisition-induced photobleaching and normalised against initial pre-bleach signal intensities. All images were processed using Nikon Elements to denoise and enhance signal intensity and contrast prior to analysis. We used ImageJ to extract Z-axis profiles for each time-lapse series to measure photobleaching and recovery curves. The resulting values were analysed in Matlab to calculate mobile fractions, which were then plotted to assess fluorescence recovery dynamics.

### HeLa cell culture

HeLa cells were cultured in DMEM, high glucose with GlutaMAX (Thermo Fisher Scientific, 61965026) supplemented with 10 % FBS (Pan Biotech, P30-3031), 1 % sodium pyruvate (Thermo Fisher Scientific, 11360039), 1 % non-essential amino acids (Thermo Fisher Scientific, 11140035) and 1 % penicillin-streptomycin (Thermo Fisher Scientific, 15140122). Cells were grown and incubated at 37°C, 5 % CO_2_ and not used over passage number 30. Two days prior to use, cells were plated at 2 x 10^4^ cells/well on poly-L-lysine (0.01 %, Sigma-Aldrich, P4707 or P6282) coated 8-well imaging chamber slides (ibidi, 80806).

### iPSC-derived Neuronal Differentiation

KOLF2.1J-WT (JIPSC001000, JAX) iPSCs^71^ with heterozygous insertion of a tetracycline-inducible human Neurogenin-2 (hNGN2) expression cassette in the *CLYBL* locus were generated as described (Casterton et al., manuscript under review). iPSCs were differentiated into cortical/forebrain patterned neurons using inducible expression of the transcription factor NGN2, method adapted from Fernanopulle *et al*.^88^. iPSCs were plated on Geltrex (Gibco, #11612149)-coated 6 well plates at a density of 5 x 10^5^ cells per well in induction media (day 1). Induction media was composed of DMEM/F-12 (Gibco, #10565018) supplemented with 1x GlutaMAX (Gibco, 35050061), 1x HEPES (Gibco, 15630080), 1x N2 (Gibco, 17502001), 1x NEAA (Gibco, 11140050) and 1x RevitaCell (Gibco, A2644501). Doxycycline (Sigma, D9891) was added at 2 µg/ml to induce expression of the NGN2 cassette. Induction media and doxycycline were refreshed on days 2 and 3 without the addition of RevitaCell. On day 4, cells were dissociated with StemPro Accutase (Gibco, 11599686) and plated at a density of 1.5 x 10^4^ cells per well on 0.1 mg/ml poly-L-ornithine (Merck, P3655) and 3-5 µg/ul laminin (Merck, L2020) coated 96 well PhenoPlates (Revvity, 6055300) in maintenance media. Maintenance media was composed of Neurobasal Plus (Gibco, A3582901) supplemented with 1 µg/ml laminin, 1x B27 Plus (Gibco, A3582801), 10 ng/ml NT-3 (Peprotech, 450-03) and 10 ng/ml BDNF (PeproTech, 450-02). This was termed DIV0 and cells were maintained with half media changes twice weekly. Import assays were performed at DIV14 or 15.

### Immunofluorescent staining and imaging of iPSC-derived neurons

iPSC-derived neurons were cultured at 37°C and 5% CO2 for 14 days on PLO (Merck, P3655) and laminin- (Merck, L2020) coated 96 well PhenoPlates (Revvity, 6055300). At DIV14, neurons were fixed in 4% paraformaldehyde (PFA) (Thermo Fisher Scientific, 11586711) for 15 mins at room temperature. PFA was then washed off three times with PBS (Thermo Fisher Scientific, 14190094) and cells permeabilised in 0.25% Triton X-100 (Thermo Fisher Scientific, A16046.AE) in PBS for 15 mins at room temperature. Cells were washed once with PBS and blocked in 10% Normal Goat Serum (NGS; Thermo Fisher Scientific, 31872) in PBS for 1 hour at room temperature. Blocking solution was removed and primary antibody (MAP-2, 1:1000 in 5% NGS in PBS, Abcam, ab92434) was added to cells and incubated at 4°C overnight. The following day, cells were washed three times with PBS. Secondary antibody (goat anti-chicken IgY Alexa FlourTM 633, 1:1000 in 5% NGS in PBS, Thermo Fisher Scientific, A21103) was added and incubated for 1 hour at room temperature in the dark. Cells were washed once with PBS and stained with DAPI (1mg/ml in PBS, Thermo Fisher Scientific, D1306) for 5 mins at room temperature in the dark then was washed once with PBS prior to storage at 4°C in PBS until imaging. Cells were imaged on an Opera Phenix high-content imaging system at 40x magnification.

### Cell Import Assays

An adaptation of a protocol from the Lowe group^43^ was employed, using digitonin to permeabilise the cells’ plasma membrane while leaving the nuclear membrane intact (semi-permeabilisation).

### HeLa Cell Permeabilisation

Two days post-plating, media on cells was replaced with DPBS and underwent half DPBS washes for 4 x 2.5 min before being replaced with permeabilisation buffer (50 mM HEPES pH 7.3, 50 mM KOAc, 8 mM MgCl_2_) for 2 minutes. Cells were then permeabilised for 10 min with permeabilisation buffer containing 50 µg/ml digitonin (Cambridge Bioscience, 2082-1), 100 µM ATP (Roche, 10519979001), 100 µM GTP (Roche, 10106399001), 4 mM creatine phosphate (Roche, 10621714001) and 20 units/ml creatine kinase (Roche, 10127566001). Cells were washed for 5 min with transport buffer (20 mM HEPES pH7.3, 110 mM KOAc, 5 mM NaOAc, 2 mM MgOAc, 2 mM DTT (Roche, 10197777001)) with 2 x 2.5 min half buffer washes. DAPI (Sigma-Aldrich, D9542), diluted to 2 µM in transport buffer, was added and incubated for at least 30 minutes prior to imaging.

### iPSC-derived Neuron Permeabilisation

Due to their sensitivity to the method used for HeLa cells, iPSC-derived neurons were permeabilised using an adapted protocol. To ensure the cells were not exposed to air during washes, 50 µl of media was left in wells and the remaining 150 µl replaced with DPBS and repeated once more. All but 50 µl of volume was removed and 150 µl permeabilisation buffer, as above, supplemented with 10% optiprep (Abcam, ab286850) was added. Half the volume was then replaced with permeabilisation buffer containing 10% optiprep and 100 µg/ml digitonin, giving a final digitonin concentration of 50 µg/ml for a 5-minute incubation. All but 50 µl volume was twice exchanged for transport buffer, as above, with the addition of 2% sucrose. 30 minutes prior to imaging, half the volume was replaced with a 2x solution of Hoechst 33342 (final dilution 1:100, Abcam, ab228551) with or without GR_20_ (final concentration of 10 µM) in transport buffer with 2% sucrose. 15 minutes prior to imaging 100 µl volume was removed and mixed with fluorescent protein. This mix was then added back to the well, giving a final fluorescent protein concentration of 2 µM. Imaging was conducted after 15 minutes.

### Cell imaging for permeabilization assays

Cells were imaged on a Nikon Eclipse Ti Inverted spinning disk confocal System with a Yokogawa CSU-X1 spinning disk unit and an Andor iXon EMMCD camera. A Plan Apochromat 40x water immersion 1.1 NA lens was used. Samples were illuminated with 405 and 488/568 nm lasers sequentially.

Live confocal microscopy was used to image the passive nuclear import of fluorescently labelled cargo from addition into the buffer into the nucleus of semi-permeabilised HeLa cells. When on the microscope, 180 µl transport buffer was removed from the well and mixed with 20 µl of fluorescent cargo before the full volume was returned to the well immediately after image acquisition began. FITC 20 kDa (Sigma-Aldrich, FD20), FITC 40 kDa (Sigma-Aldrich, FD40), FITC 70 kDa (Sigma-Aldrich, 90718) and Alexa Fluor 10kDa (647, D22914) dextrans were used at a final assay concentration of 10 µg/ml. GFP mutants were used at a final assay concentration of 2 µM, except highly hydrophobic mutants efGFP 8W and 8L which were used at 0.2 µM to allow quantification of nuclear import in addition to aggregation. Images were taken every 2.5 seconds for 4 minutes. For iPSC-derived neurons, confocal microscopy was used to assess passive nuclear import 15 minutes post addition of cargo, using the same imaging as for HeLa cells but only capturing the endpoint to increase throughput.

### U2OS cell culture

U2OS-CRISPR-NUP96-mEGFP cells (Cytion, #300174) were cultured in McCoy’s 5A (Modified) Medium (Thermo Fisher Scientific, 26600023) supplemented with 10% fetal bovine serum (Pan Biotech, P30-3031), 1% non-essential amino acids (Thermo Fisher Scientific, 11140035), 1% penicillin/streptomycin (Thermo Fisher Scientific, 15140122), and 1% sodium pyruvate (Thermo Fisher Scientific, 11360039). Cells were maintained at 37°C in a humidified incubator with 5% CO2. For imaging, cells were seeded onto µ-Slide glass bottomed 8-well chamber slides (Ibidi, 80807) pre-coated with 2M glycine (VMR chemicals, #444495D) for 30-60 minutes to promote optimal adhesion and reduce fluorescence background for imaging. Cells were incubated with 100 nM ATTO655 or GR_20_-ATTO655 for 2 hours in growth media. For live-cell imaging of nuclear accumulation cells underwent a single wash step in media prior to imaging. For single molecule localisation GR_20_ and NPC analyses, cells continued into preparation for DNA-PAINT.

### DNA-PAINT

Immunostaining with DNA-PAINT (Points Accumulation for Imaging in Nanoscale Topography) was carried out to improve the spatial resolution of GFP, following the protocol adapted from Schlichthaerle *et al.,* (2019)^89^. Cells were rinsed once with PBS, followed by a 20-second wash with 2.4 % paraformaldehyde (PFA, Thermo Fisher Scientific, 11586711) in PBS. Permeabilization was performed using 0.4 % Triton X-100 in PBS for 10 seconds. Cells were then fixed again in 2.4 % PFA for 30 minutes, washed thoroughly three times with PBS, and quenched with 50 mM ammonium chloride (Sigma-Aldrich, A5354) for 4 minutes. This step was followed by three 5-minute PBS washes. Next, cells were blocked with antibody incubation buffer in PBS for 30 minutes at room temperature to minimise nonspecific binding. For specific labelling of Nup96-GFP, single-domain anti-GFP antibody with FAST docking strand was diluted (1:2000) in antibody buffer and incubated overnight at 4°C. After incubation, cells were washed three times with washing buffer. For imaging preparation, a final wash with imaging buffer containing 2 nM Cy3B imager, which binds to the FAST docking strand of the anti-GFP nanobody, and fiducial markers (Thermo Fisher Scientific, T7279) were applied. Anti-GFP nanobody and imaging kit including buffers is available from Massive Photonics (MASSIVE-TAG-Q anti-GFP Fluorescent Protein and Tag Imaging Kit).

### Imaging of polyGR and NPCs in U2OS-NUP96-GFP cells

A Nikon Eclipse TiE inverted fluorescence microscope equipped with high-NA oil immersion objectives (Nikon CFI Apo TIRF 100XC Oil NA1.49 and Nikon CFI SR HP Apo TIRF 100XAC Oil NA1.49), a TIRF (Total Internal Reflection Fluorescence) illuminator, and a high-sensitivity camera (Andor iXon EMCCD) was used. First, a semi-bleaching process was performed to decrease the background signal originating from the bottom of the coverslip, ensuring enhanced signal-to-noise ratio. For live-cell imaging of nuclear accumulation, the TIRF angle was adjusted to 65° to enable selective illumination of NPCs and ATTO655 dye molecules near the nuclear membrane. For single molecule localisation GR_20_ and NPC analyses, STORM (Stochastic Optical Reconstruction Microscopy) with TIRF was used. We examined the U2OS-NUP96-GFP cells using a GFP nanobody conjugated to Cy3B imager strands alongside fluorescently labelled DPRs (GR_20_-ATTO655) or ATTO655 dye alone. By integrating multi-colour STORM with TIRF, we achieved a lateral resolution of approximately 20 nm in relation to NPCs, enabling the visualisation of individual Nup96 protein and ATTO655-labelled molecules with high precision. Imaging of Cy3B and ATTO655 was performed sequentially to prevent spectral overlap and ensure accurate localisation. 10,000 frames were acquired, and localisations were computationally reconstructed into a high-resolution image. ThunderSTORM ImageJ plugin was used for image processing, including multi-colour alignment via fiducial markers and drift correction to enhance structural clarity. To generate representative NPC images, particle fusion was applied to align and average multiple individual NPCs, improving signal-to-noise ratio and revealing conserved nanoscale features^90, 91^.

### Disorder Analysis

Intrinsic disorder across the Nup98 sequence was predicted using metapredict v3^92^ (metapredict-IDP model; default parameters). The Nup98 amino-acid sequence (residues 1–880 for Nup98 from autolysis of precursor Nup98-Nup96, UniProt P52948) was used as input, and per-residue disorder scores were generated. Regions with disorder scores > 0.5 were classified as intrinsically disordered.

### Alpha fold

To assess whether the introduced mutations were predicted to alter protein structure, we generated structural models for all GFP variants using AlphaFold2, providing each variant’s exact amino acid sequence as input. The resulting predicted structures were examined for any mutation-associated conformational changes. Structural models were then imported into the RCSB Protein Data Bank (PDB) visualization tools for inspection and comparison across variants.

### Coarse-grained simulations

All coarse-grained simulations utilized the PIMMS simulation engine (https://github.com/holehouse-lab/PIMMS/). PIMMS is a lattice-based coarse-grained simulation engine used previously to characterize the phase behaviour of low-complexity disordered domains^93–95^, one-bead-per-residue resolution model parameterized by optimizing pairwise interactions to recapitulate dimensions of disordered proteins measured using small-angle X-ray scattering (SAXS). The specific parameter file, as well as example key files to run the simulations, can be found at https://github.com/holehouse-lab/supportingdata/tree/master/2026/solomon_et_al_2026.

For all simulations, move parameters were as follows: CRANKSHAFT_STEPS: 1,000,000; CRANKSHAFT_MODE: PROPORTIONAL; MOVE_CRANKSHAFT: 0.9; MOVE_CHAIN_TRANSLATE: 0.1, and all other move types set to 0. Further, simulations were run with HARDWALL: True and with TEMPERATURE: 50. We note that this temperature (defined based on force field parameterization) is not directly interpretable in Fahrenheit or Celsius but rather is a parameter that impacts the energy in the simulation.

Simulations were run in a multi-stage format to ensure robust equilibration, with an initial Nup98 equilibration simulation followed by a partitioning simulation.

Nup98 equilibration simulations were run with 100 copies of a Nup98 FG fragment (residues 2-151) in a [100x100x100] box (dimensions in lattice units). Chains were initialized within a central [29x29x29] box using a 1-step initial phase, followed by 10,000 steps in the full-size simulation box, enabling a single Nup98 condensate to form and equilibrate. We note that running simulations without this procedure eventually leads to a single Nup98 condensate; this pre-initialization acts to aid convergence and ensure a single condensate can form in all independent simulations. These equilibrated Nup98 condensates were then used as starting points for partitioning simulations.

For partitioning simulations, we took equilibrated Nup98 condensate configurations generated using the Nup98 equilibration simulations and used PIMMS’ restart capabilities to initialize a new simulation where additional chains were added around the Nup98 condensate. Specifically, for all simulations where a probe or 20-mer GR_10_ peptide was added, the following parameters were used: RESTART_FILE: nup_1_153_eq_short.pimms; EXPERIMENTAL_FEATURES: True (Note: this allowed us to add in additional chains not original in the equilibration simulation); DIMENSIONS: 115 115 115, N_STEPS: 20,000; EQUILIBRATION: 1,000; XTC_FREQ: 40. In addition, either probe chains or probe chains + GR_20_ chains were included in the partitioning simulations. The first 14,000 steps were discarded for equilibration, leaving the final 6,000 steps (150 frames) for simulation analysis. Every simulation ran two independent replicates. Partitioning was quantified by the extent with which peptides were incorporated into the Nup98 condensate, based on the CLUSTER.dat output file and the associated cluster membership files.

### Statistics

All statistical testing, as detailed in the figure legends, was performed using GraphPad Prism 10.5.0. Condensate and cell data are presented as mean ± SEM, simulation data as mean ± SD. Statistical comparisons between two groups were performed using t-tests, while comparisons among more than two groups were conducted using ANOVAs with Tukey’s multiple comparisons testing. A significance threshold of p < 0.05 was applied; p values <0.1 are also stated in figures to show trends. Additional details regarding the statistical analyses and N numbers are provided in the figure legends. All images are representative from at least three independent experiments, unless otherwise stated.

## Acknowledgements

We thank Prof. Dirk Gorlich and Dr. Sheung Chun Ng for sharing plasmids for the efGFP mutant series and associated proteases required for recombinant purification and providing recombinant SinGFP4a and NTF2-Alexa488. We also thank Dr. Sheung Chun Ng for thoughtful suggestions related to the FG phase assay experiments and comments on the manuscript. We thank George Chennel and Chen Liang from the Wohl Cellular Imaging Centre, (King’s College London, London, UK) for training and technical support with microscopy. This project was predominantly funded by a UK Dementia Research Institute award to S.M (UK DRI-6203) through UK DRI Ltd, principally funded by the Medical Research Council. D.A.S. is supported by The Lady Edith Wolfson Foundation Fellowship Programme fellowship from the Motor Neurone Association (MNDA) (Solomon/Oct23/2329-799). This work as also funded by HFSP grant RGP0015/2022 to A.S.H.. iPSC line development was funded by a MNDA PhD studentship and UK Dementia Research Institute awards to MD.R (Ruepp/Oct22/911-792 and UK DRI-6204).

## Author Contributions

S.M., D.A.S., H.B.S., A.S.H., designed and conceptualised this study and S.M. secured funding. S.M., and D.A.S., oversaw project administration. S.M., A.S.H., H.B.S., D.A.S., provided supervision. D.A.S., R.J.E., MT.SK., S.M.K., O.H., E.W., S.L., I.L.C., S.M., performed the experiments and analysed the data. D.A.S., and S.M.K., performed all recombinant protein purifications. D.A.S., designed, performed and analysed the FG phase assays. R.J.E., designed, performed and analysed the coarse-grained simulations. MT.SK., S.M., O.H, E.W., D.A.S., designed, performed and analysed permeabilised cell assays in HeLa cells. MT.SK, O.H., D.A.S., S.M, designed, performed and analysed permeabilised cell assays in iPSC neurons. S.L., D.A.S., S.M., designed, performed and analysed single molecule localisation experiments of nuclear pore localisation. N.O.B., J.A., and MD.R. provided resources prior to publication. D.A.S., R.J.E., S.L., A.S.H., and S.M designed and prepared visualisations. D.A.S., and R.J.E., wrote the original draft. D.A.S., R.J.E., S.M., A.S.H., H.B.S, J.A., MD.R., reviewed, edited and wrote revised drafts. All authors approved the final version of the manuscript.

## Ethics declarations

N/A

## References

1. Beck, M. & Hurt, E. The nuclear pore complex: understanding its function through structural insight. Nature Reviews Molecular Cell Biology 18, 73–89 (2017).

2. Schmidt, H.B. & Görlich, D. Transport Selectivity of Nuclear Pores, Phase Separation, and Membraneless Organelles. Trends in Biochemical Sciences 41, 46–61 (2016).

3. Yang, Y. et al. Nuclear transport proteins: structure, function, and disease relevance. Signal Transduction and Targeted Therapy 8, 425 (2023).

4. Coyne, A.N. & Rothstein, J.D. Nuclear pore complexes — a doorway to neural injury in neurodegeneration. Nature Reviews Neurology 18, 348–362 (2022).

5. Fare, C.M. & Rothstein, J.D. Nuclear pore dysfunction and disease: a complex opportunity. Nucleus 15, 2314297 (2024).

6. Cristi, A.C., Rapuri, S. & Coyne, A.N. Nuclear pore complex and nucleocytoplasmic transport disruption in neurodegeneration. FEBS Letters 597, 2546–2566 (2023).

7. Lusk, C.P. & King, M.C. The nucleus: keeping it together by keeping it apart. Current Opinion in Cell Biology 44, 44–50 (2017).

8. Spead, O., Zaepfel, B.L. & Rothstein, J.D. Nuclear Pore Dysfunction in Neurodegeneration. Neurotherapeutics 19, 1050–1060 (2022).

9. Ling, S.-C., Polymenidou, M. & Cleveland, D.W. Converging Mechanisms in ALS and FTD: Disrupted RNA and Protein Homeostasis. Neuron 79, 416–438 (2013).

10. Renton, A.E. et al. A Hexanucleotide Repeat Expansion in *C9ORF72* Is the Cause of Chromosome 9p21-Linked ALS-FTD. Neuron 72, 257–268 (2011).

11. DeJesus-Hernandez, M. et al. Expanded GGGGCC Hexanucleotide Repeat in Noncoding Region of *C9ORF72* Causes Chromosome 9p-Linked FTD and ALS. Neuron 72, 245–256 (2011).

12. Mackenzie, I.R. et al. Dipeptide repeat protein pathology in C9ORF72 mutation cases: clinico-pathological correlations. Acta Neuropathologica 126, 859–879 (2013).

13. Mackenzie, I.R. & Neumann, M. Subcortical TDP-43 pathology patterns validate cortical FTLD-TDP subtypes and demonstrate unique aspects of C9orf72 mutation cases. Acta Neuropathologica 139, 83–98 (2020).

14. Mori, K. et al. Bidirectional transcripts of the expanded C9orf72 hexanucleotide repeat are translated into aggregating dipeptide repeat proteins. Acta Neuropathologica 126, 881–893 (2013).

15. Ash, P.E.A. et al. Unconventional Translation of *C9ORF72* GGGGCC Expansion Generates Insoluble Polypeptides Specific to c9FTD/ALS. Neuron 77, 639–646 (2013).

16. Mizielinska, S. et al. C9orf72 repeat expansions cause neurodegeneration in Drosophila through arginine-rich proteins. Science 345, 1192–1194 (2014).

17. Kwon, I. et al. Poly-dipeptides encoded by the C9orf72 repeats bind nucleoli, impede RNA biogenesis, and kill cells. Science 345, 1139–1145 (2014).

18. Wen, X., et al. Antisense Proline-Arginine RAN Dipeptides Linked to C9ORF72-ALS/FTD Form Toxic Nuclear Aggregates that Initiate In Vitro and In Vivo Neuronal Death. Neuron 84, 1213–1225 (2014).

19. McGoldrick, P. & Robertson, J. Unraveling the impact of disrupted nucleocytoplasmic transport systems in C9orf72-associated ALS. Frontiers in Cellular Neuroscience 17 (2023).

20. Zhang, K. et al. The C9orf72 repeat expansion disrupts nucleocytoplasmic transport. Nature 525, 56–61 (2015).

21. Freibaum, B.D. et al. GGGGCC repeat expansion in C9ORF72 compromises nucleocytoplasmic transport. Nature 525, 129–133 (2015).

22. Jovičić, A. et al. Modifiers of C9orf72 dipeptide repeat toxicity connect nucleocytoplasmic transport defects to FTD/ALS. Nature Neuroscience 18, 1226–1229 (2015).

23. Boeynaems, S. et al. Drosophila screen connects nuclear transport genes to DPR pathology in c9ALS/FTD. Scientific Reports 6, 20877 (2016).

24. Solomon, D.A. et al. A feedback loop between dipeptide-repeat protein, TDP-43 and karyopherin-α mediates C9orf72-related neurodegeneration. Brain 141, 2908–2924 (2018).

25. Hutten, S. et al. Nuclear Import Receptors Directly Bind to Arginine-Rich Dipeptide Repeat Proteins and Suppress Their Pathological Interactions. Cell Reports 33, 108538 (2020).

26. Hayes, L.R., Duan, L., Bowen, K., Kalab, P. & Rothstein, J.D. C9orf72 arginine-rich dipeptide repeat proteins disrupt karyopherin-mediated nuclear import. eLife 9, e51685 (2020).

27. Ryan, S., Rollinson, S., Hobbs, E. & Pickering-Brown, S. C9orf72 dipeptides disrupt the nucleocytoplasmic transport machinery and cause TDP-43 mislocalisation to the cytoplasm. Scientific Reports 12, 4799 (2022).

28. Lee, K.-H. et al. C9orf72 Dipeptide Repeats Impair the Assembly, Dynamics, and Function of Membrane-Less Organelles. Cell 167, 774–788.e717 (2016).

29. Lin, Y. et al. Toxic PR Poly-Dipeptides Encoded by the C9orf72 Repeat Expansion Target LC Domain Polymers. Cell 167, 789–802.e712 (2016).

30. Borcherds, W., Bremer, A., Borgia, M.B. & Mittag, T. How do intrinsically disordered protein regions encode a driving force for liquid–liquid phase separation? Current Opinion in Structural Biology 67, 41–50 (2021).

31. Banani, S.F., Lee, H.O., Hyman, A.A. & Rosen, M.K. Biomolecular condensates: organizers of cellular biochemistry. Nature Reviews Molecular Cell Biology 18, 285–298 (2017).

32. Holehouse, A.S. & Kragelund, B.B. The molecular basis for cellular function of intrinsically disordered protein regions. Nature Reviews Molecular Cell Biology 25, 187–211 (2024).

33. Banerjee, P.R. et al. Dissecting the biophysics and biology of intrinsically disordered proteins. Trends in Biochemical Sciences 49, 101–104 (2024).

34. Mathieu, C., Pappu, R.V. & Taylor, J.P. Beyond aggregation: Pathological phase transitions in neurodegenerative disease. Science 370, 56–60 (2020).

35. Sabari, B.R., Dall’Agnese, A. & Young, R.A. Biomolecular Condensates in the Nucleus. Trends in Biochemical Sciences 45, 961–977 (2020).

36. Ng, S.C. et al. Barrier properties of Nup98 FG phases ruled by FG motif identity and inter-FG spacer length. Nature Communications 14, 747 (2023).

37. Schmidt, H.B. & Görlich, D. Nup98 FG domains from diverse species spontaneously phase-separate into particles with nuclear pore-like permselectivity. eLife 4, e04251 (2015).

38. Hülsmann, B.B., Labokha, A.A. & Görlich, D. The permeability of reconstituted nuclear pores provides direct evidence for the selective phase model. Cell 150, 738–751 (2012).

39. Fu, L. et al. Nuclear pore passage of the HIV capsid is driven by its unusual surface amino acid composition. Nat Struct Mol Biol 32, 2476–2491 (2025).

40. Mohr, D., Frey, S., Fischer, T., Güttler, T. & Görlich, D. Characterisation of the passive permeability barrier of nuclear pore complexes. The EMBO journal 28, 2541–2553 (2009).

41. Frey, S. et al. Surface Properties Determining Passage Rates of Proteins through Nuclear Pores. Cell 174, 202–217.e209 (2018).

42. Naim, B. et al. Passive and facilitated transport in nuclear pore complexes is largely uncoupled. J Biol Chem 282, 3881–3888 (2007).

43. Lowe, A.R. et al. Importin-β modulates the permeability of the nuclear pore complex in a Ran-dependent manner. eLife 4, e04052 (2015).

44. Andersen, K.R. et al. Scaffold nucleoporins Nup188 and Nup192 share structural and functional properties with nuclear transport receptors. eLife 2, e00745 (2013).

45. Kozai, T. et al. Karyopherins remodel the dynamic organization of the nuclear pore complex transport barrier. Nature Cell Biology, 1–13 (2025).

46. Frey, S., Richter, R.P. & Görlich, D. FG-Rich Repeats of Nuclear Pore Proteins Form a Three-Dimensional Meshwork with Hydrogel-Like Properties. Science 314, 815–817 (2006).

47. Frey, S. & Görlich, D. A Saturated FG-Repeat Hydrogel Can Reproduce the Permeability Properties of Nuclear Pore Complexes. Cell 130, 512–523 (2007).

48. Milles, S. & Lemke, E.A. Single Molecule Study of the Intrinsically Disordered FG-Repeat Nucleoporin 153. Biophysical Journal 101, 1710–1719 (2011).

49. Labokha, A.A. et al. Systematic analysis of barrier-forming FG hydrogels from Xenopus nuclear pore complexes. The EMBO Journal 32, 204–218 (2013).

50. Ng, S.C., Güttler, T. & Görlich, D. Recapitulation of selective nuclear import and export with a perfectly repeated 12mer GLFG peptide. Nature Communications 12, 4047 (2021).

51. Najbauer, E.E., Ng, S.C., Griesinger, C., Görlich, D. & Andreas, L.B. Atomic resolution dynamics of cohesive interactions in phase-separated Nup98 FG domains. Nature Communications 13, 1494 (2022).

52. Frey, S. & Görlich, D. FG/FxFG as well as GLFG repeats form a selective permeability barrier with self-healing properties. The EMBO Journal 28, 2554–2567 (2009).

53. Konishi, H.A. & Yoshimura, S.H. Interactions between non-structured domains of FG-and non-FG-nucleoporins coordinate the ordered assembly of the nuclear pore complex in mitosis. Faseb j 34, 1532–1545 (2020).

54. Ng, S.C. & Görlich, D. A simple thermodynamic description of phase separation of Nup98 FG domains. Nature Communications 13, 6172 (2022).

55. Yu, M. et al. Visualizing the disordered nuclear transport machinery in situ. Nature 617, 162–169 (2023).

56. Fu, L. et al. HIV-1 capsids enter the FG phase of nuclear pores like a transport receptor. Nature 626, 843–851 (2024).

57. Dickson, C.F. et al. The HIV capsid mimics karyopherin engagement of FG-nucleoporins. Nature 626, 836–842 (2024).

58. Eftekharzadeh, B. et al. Tau Protein Disrupts Nucleocytoplasmic Transport in Alzheimer’s Disease. Neuron 99, 925–940.e927 (2018).

59. Lin, Y.-C. et al. Interactions between ALS-linked FUS and nucleoporins are associated with defects in the nucleocytoplasmic transport pathway. Nature neuroscience 24, 1077–1088 (2021).

60. Diez, L. et al. Phosphorylation but Not Oligomerization Drives the Accumulation of Tau with Nucleoporin Nup98. International Journal of Molecular Sciences 23, 3495 (2022).

61. Gleixner, A.M. et al. NUP62 localizes to ALS/FTLD pathological assemblies and contributes to TDP-43 insolubility. Nature Communications 13, 3380 (2022).

62. Dickson, J.R., Frosch, M.P. & Hyman, B.T. Altered localization of nucleoporin 98 in primary tauopathies. Brain Communications 5, fcac334 (2023).

63. Dubey, S.K. et al. Aberrant nuclear pore complex degradation contributes to neurodegeneration in VCP disease. Neuron (2025).

64. Zhang, K. et al. Stress Granule Assembly Disrupts Nucleocytoplasmic Transport. Cell 173, 958–971.e917 (2018).

65. Park, J. et al. Poly(GR) interacts with key stress granule factors promoting its assembly into cytoplasmic inclusions. Cell Reports 42, 112822 (2023).

66. Van Nerom, M. et al. C9orf72-linked arginine-rich dipeptide repeats aggravate pathological phase separation of G3BP1. Proceedings of the National Academy of Sciences 121, e2402847121 (2024).

67. Zhang, Y.-J. et al. Poly(GR) impairs protein translation and stress granule dynamics in C9orf72-associated frontotemporal dementia and amyotrophic lateral sclerosis. Nature Medicine 24, 1136–1142 (2018).

68. Miyagi, T. et al. Differential toxicity and localization of arginine-rich *C9ORF72* dipeptide repeat proteins depend on de-clustering of positive charges. iScience 26, 106957 (2023).

69. Solomon, D.A., Smikle, R., Reid, M.J. & Mizielinska, S. Altered Phase Separation and Cellular Impact in C9orf72-Linked ALS/FTD. Frontiers in Cellular Neuroscience 15 (2021).

70. Holehouse, A.S. & Alberti, S. Molecular determinants of condensate composition. Mol Cell 85, 290–308 (2025).

71. Pantazis, C.B. et al. A reference human induced pluripotent stem cell line for large-scale collaborative studies. Cell Stem Cell 29, 1685–1702.e1622 (2022).

72. Shi, K.Y. et al. Toxic PRn poly-dipeptides encoded by the C9orf72 repeat expansion block nuclear import and export. Proceedings of the National Academy of Sciences of the United States of America 114, E1111–E1117 (2017).

73. Ribbeck, K. & Görlich, D. The permeability barrier of nuclear pore complexes appears to operate via hydrophobic exclusion. The EMBO Journal 21, 2664–2671 (2002).

74. Villegas, J.A. & Levy, E.D. A unified statistical potential reveals that amino acid stickiness governs nonspecific recruitment of client proteins into condensates. Protein Science 31, e4361 (2022).

75. Jafarinia, H., van der Giessen, E. & Onck, P.R. C9orf72 polyPR directly binds to various nuclear transport components. eLife 12, RP89694 (2024).

76. Vanneste, J. et al. C9orf72-generated poly-GR and poly-PR do not directly interfere with nucleocytoplasmic transport. Scientific Reports 9, 15728 (2019).

77. Ederle, H. et al. Nuclear egress of TDP-43 and FUS occurs independently of Exportin-1/CRM1. Scientific Reports 8, 7084 (2018).

78. Duan, L. et al. Nuclear RNA binding regulates TDP-43 nuclear localization and passive nuclear export. Cell Reports 40, 111106 (2022).

79. Junod, S.L., Kelich, J.M., Ma, J. & Yang, W. Nucleocytoplasmic transport of intrinsically disordered proteins studied by high-speed super-resolution microscopy. Protein Science: A Publication of the Protein Society 29, 1459–1472 (2020).

80. Yu, W., Rush, C., Tingey, M., Junod, S. & Yang, W. Application of Super-resolution SPEED Microscopy in the Study of Cellular Dynamics. Chemical & Biomedical Imaging 1, 356–371 (2023).

81. Coyne, A.N. et al. G4C2 Repeat RNA Initiates a POM121-Mediated Reduction in Specific Nucleoporins in C9orf72 ALS/FTD. Neuron 107, 1124–1140.e1111 (2020).

82. Coyne, A.N. et al. Nuclear accumulation of CHMP7 initiates nuclear pore complex injury and subsequent TDP-43 dysfunction in sporadic and familial ALS. Science Translational Medicine 13, eabe1923 (2021).

83. Frey, S. & Görlich, D. A new set of highly efficient, tag-cleaving proteases for purifying recombinant proteins. Journal of Chromatography A 1337, 95–105 (2014).

84. Contreras-Martos, S. et al. Quantification of Intrinsically Disordered Proteins: A Problem Not Fully Appreciated. Frontiers in Molecular Biosciences Volume 5 - 2018 (2018).

85. Frey, S. & Görlich, D. Purification of protein complexes of defined subunit stoichiometry using a set of orthogonal, tag-cleaving proteases. Journal of Chromatography A 1337, 106–115 (2014).

86. Raran-Kurussi, S., Cherry, S., Zhang, D. & Waugh, D.S. Removal of Affinity Tags with TEV Protease, in Heterologous Gene Expression in E.coli: Methods and Protocols. (ed. N.A. Burgess-Brown) 221–230 (Springer, New York, NY; 2017).

87. Palmer, I. & Wingfield, P.T. Preparation and extraction of insoluble (inclusion-body) proteins from Escherichia coli. Current Protocols in Protein Science Chapter 6, 6.3.1–6.3.20 (2012).

88. Fernandopulle, M.S. et al. Transcription Factor-Mediated Differentiation of Human iPSCs into Neurons. Curr Protoc Cell Biol 79, e51 (2018).

89. Schlichthaerle, T. et al. Direct Visualization of Single Nuclear Pore Complex Proteins Using Genetically-Encoded Probes for DNA-PAINT. Angew Chem Int Ed Engl 58, 13004–13008 (2019).

90. Heydarian, H. et al. Template-free 2D particle fusion in localization microscopy. Nat Methods 15, 781–784 (2018).

91. Wang, W. et al. Particle fusion of super-resolution data reveals the unit structure of Nup96 in Nuclear Pore Complex. Sci Rep 13, 13327 (2023).

92. Emenecker, R.J., Griffith, D. & Holehouse, A.S. Metapredict: a fast, accurate, and easy-to-use predictor of consensus disorder and structure. Biophys J 120, 4312–4319 (2021).

93. Holehouse, A.S. & Pappu, R.V., Edn. 0.24 pre-beta (Zenodo, 2019).

94. Martin, E.W. et al. Valence and patterning of aromatic residues determine the phase behavior of prion-like domains. Science 367, 694–699 (2020).

95. Holehouse, A.S., Ginell, G.M., Griffith, D. & Böke, E. Clustering of Aromatic Residues in Prion-like Domains Can Tune the Formation, State, and Organization of Biomolecular Condensates. Biochemistry 60, 3566–3581 (2021).

