## Supplementary Figures for "C9ORF72-derived polyGR polypeptides disrupt passive nucleocytoplasmic transport by tuning protein affinity for the nuclear pore barrier"

### Extended Data Figures

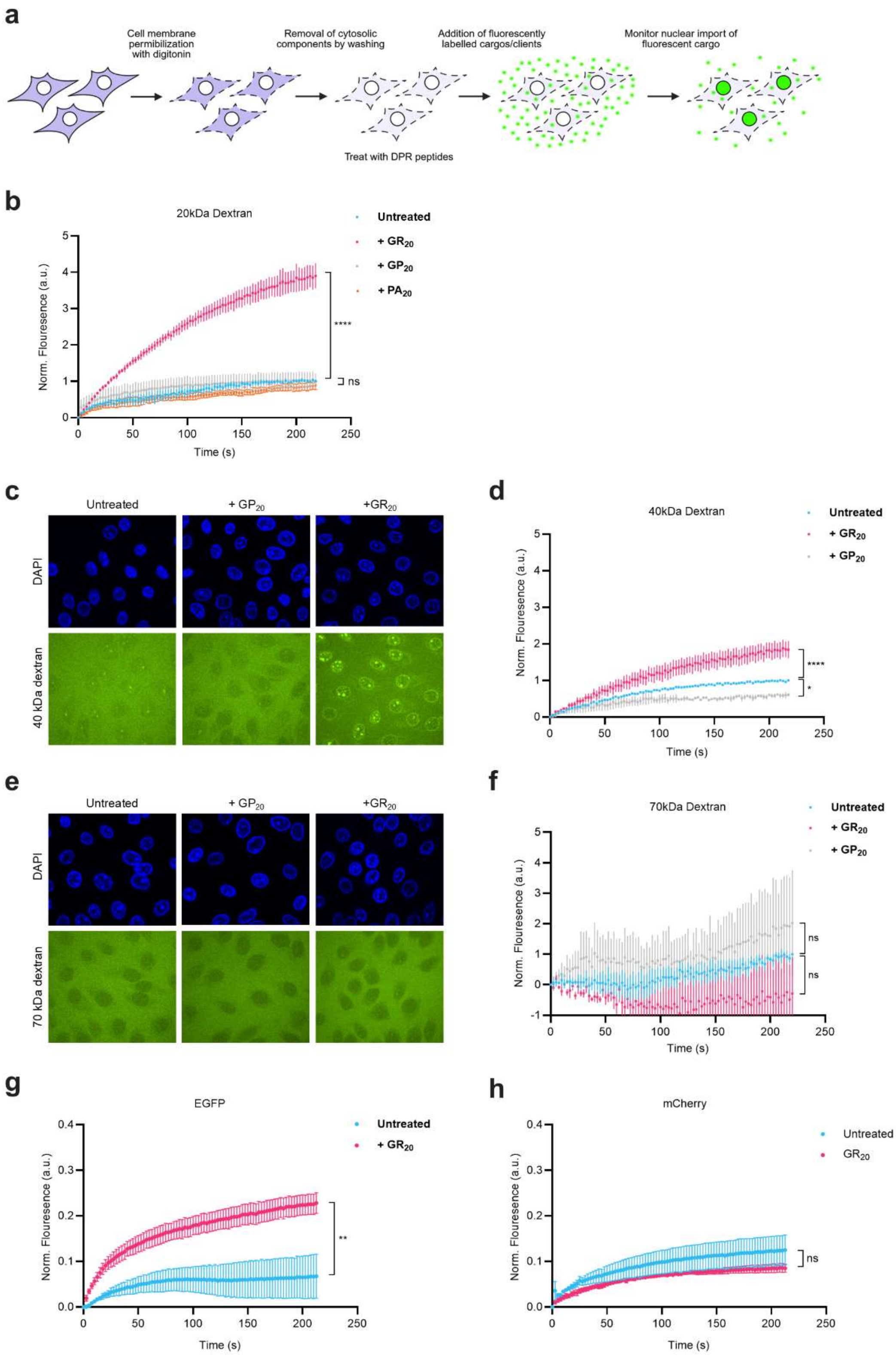


**Extended Data Figure 1 | PolyGR enhances passive nuclear import of dextran clients in a size dependent manner but exhibits client-specific effects on similarly sized proteins. a,** Schematic of the permeabilised-cell nuclear import assay used to assess passive transport of fluorescent dextran clients in HeLa cells. **b,** Passive nuclear import time course of 20 kDa dextran following treatment with $GR_{20}$, $GP_{20}$, or $PA_{20}$. PolyGR markedly increased nuclear accumulation of the 20 kDa client, whereas polyGP and polyPA had no effect (two-way ANOVA with Dunnett's multiple comparisons test; n = 7 biological replicates for untreated, 3 for polyGR and 2 for polyGP and polyPA; ****p < 0.0001; ns, not significant). **c, e, f,** PolyGR caused only a mild enhancement of 40 kDa dextran influx and no detectable change in 70 kDa dextran transport, and neither polyGP nor polyPA altered import of either client (two-way ANOVA with Dunnett's multiple comparisons test; n = 5 biological replicates for untreated, 3 for polyGR and 2 for polyGP; ****p < 0.0001; *p < 0.01; ns, not significant). **g,h,** Time courses of nuclear import of EGFP and mCherry clients, untreated and in the presence of polyGR. Despite being of comparable size, polyGR selectively enhanced nuclear import of EGFP but not mCherry, indicating that size alone is insufficient to explain polyGR-mediated modulation of passive nuclear transport and that client-specific properties contribute to transport outcomes. Scale bars = 20 μm.

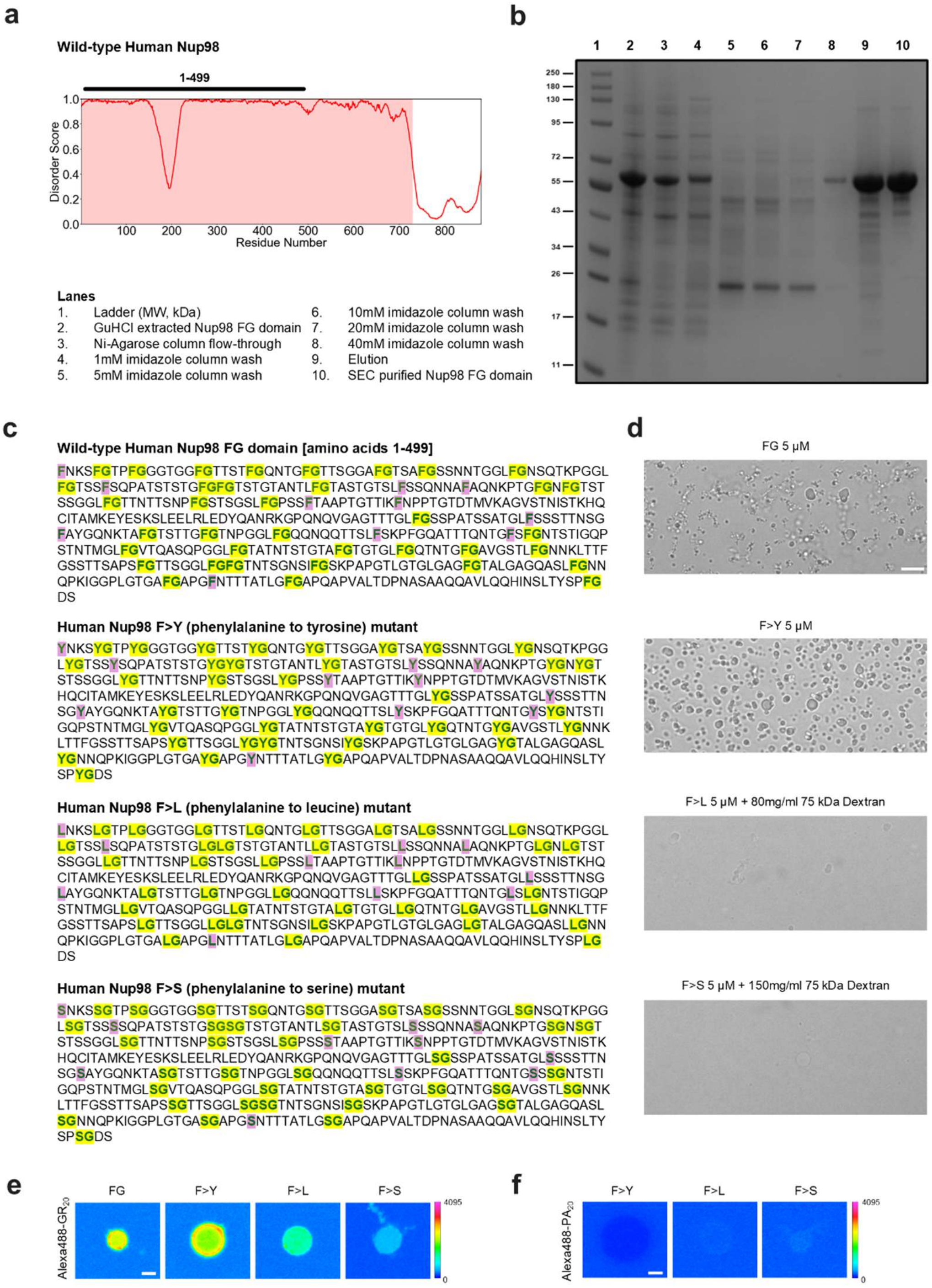


**Extended Data Figure 2 | Aromatic and hydrophobic interactions determine Nup98 FG phase separation and polyGR partitioning. a,** Plot of intrinsic disorder across human Nup98 protein, with regions of disorder (score>0.5) highlighted and 1-499 FG domain annotated. **b**, Coomassie stain of recombinant protein purification of the human Nup98 FG domain (amino acids 1–499). **c,** Design of Nup98 FG variants used to dissect aromatic and hydrophobic contributions to polyGR interactions: F>Y (preserves aromaticity and hydrophobicity), F>L (retains hydrophobicity but removes aromaticity), and F>S (lacks both aromatic and hydrophobic interactions). **d,** Phase separation behaviour of wild-type and variant FG domains upon dilution from denaturant. Wild-type and F>Y readily form condensates, F>L requires molecular crowding to phase separate, and F>S displays markedly reduced phase separation, forming only sparse condensates at high crowder concentrations. Partitioning of (Alexa488)-labelled $GR_{20}$ (**e**) and $PA_{20}$ (**f**) into FG condensates of each variant. PolyGR partitioning is strongest in wild-type and F>Y condensates, reduced in F>L, and weakest in F>S, whereas polyPA is excluded from wild-type and F>Y condensates and shows only weak entry into the more permeable F>L and F>S condensates. These results indicate that polyGR engages nuclear pore FG domains primarily through aromatic interactions, with hydrophobicity providing additional contributions. Scale bars = 2 μm.

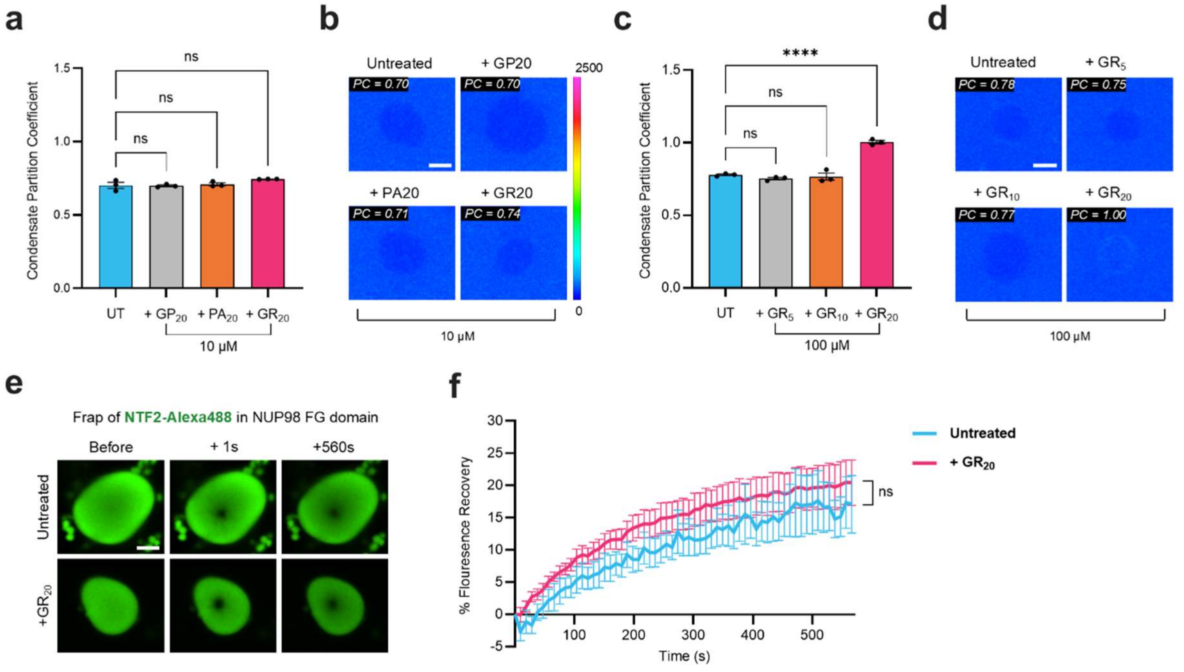


**Extended Data Figure 3 | PolyGR tunes FG-phase entry in a concentration- and length-dependent manner without altering condensate dynamics. a,b,** Partitioning of EGFP into Nup98 FG condensates is unchanged in the presence of either 10 μM polyGR, polyGP, or polyPA compared to untreated (UT). Images show representative condensate heatmap intensity scaling together with corresponding condensate partition coefficients (one-way ANOVA with Tukey's multiple comparisons test; n = 5 biological replicates, ≥5 condensates per replicate; ****p < 0.0001; ns, not significant). **c,d,** PolyGR peptide length modulated EGFP influx into the FG phase, with a significant increase relative to UT observed only for GR$_{20}$, but not for GR$_{5}$ or GR$_{10}$ (all 100 μM), demonstrating that interaction strength scales with peptide length. Images show representative condensate heatmap intensity scaling together with corresponding condensate partition coefficients (one-way ANOVA with Tukey's multiple comparisons test; n = 5 biological replicates, ≥5 condensates per replicate; ****p < 0.0001; ns, not significant). **e,f,** Fluorescence recovery after photobleaching (FRAP) of NTf2 within FG condensates revealed unchanged NTf2 mobility in the presence of 100 μM GR$_{20}$, indicating that polyGR alters FG-phase selectivity without affecting condensate dynamics. Mean fluorescence recovery curves are shown for untreated (UT) and polyGR-treated condensates, with recovery rate constants (K) fitted for each replicate (Mann–Whitney U test; n = 4 biological replicates; ns, not significant). Scale bars = 2 μm.

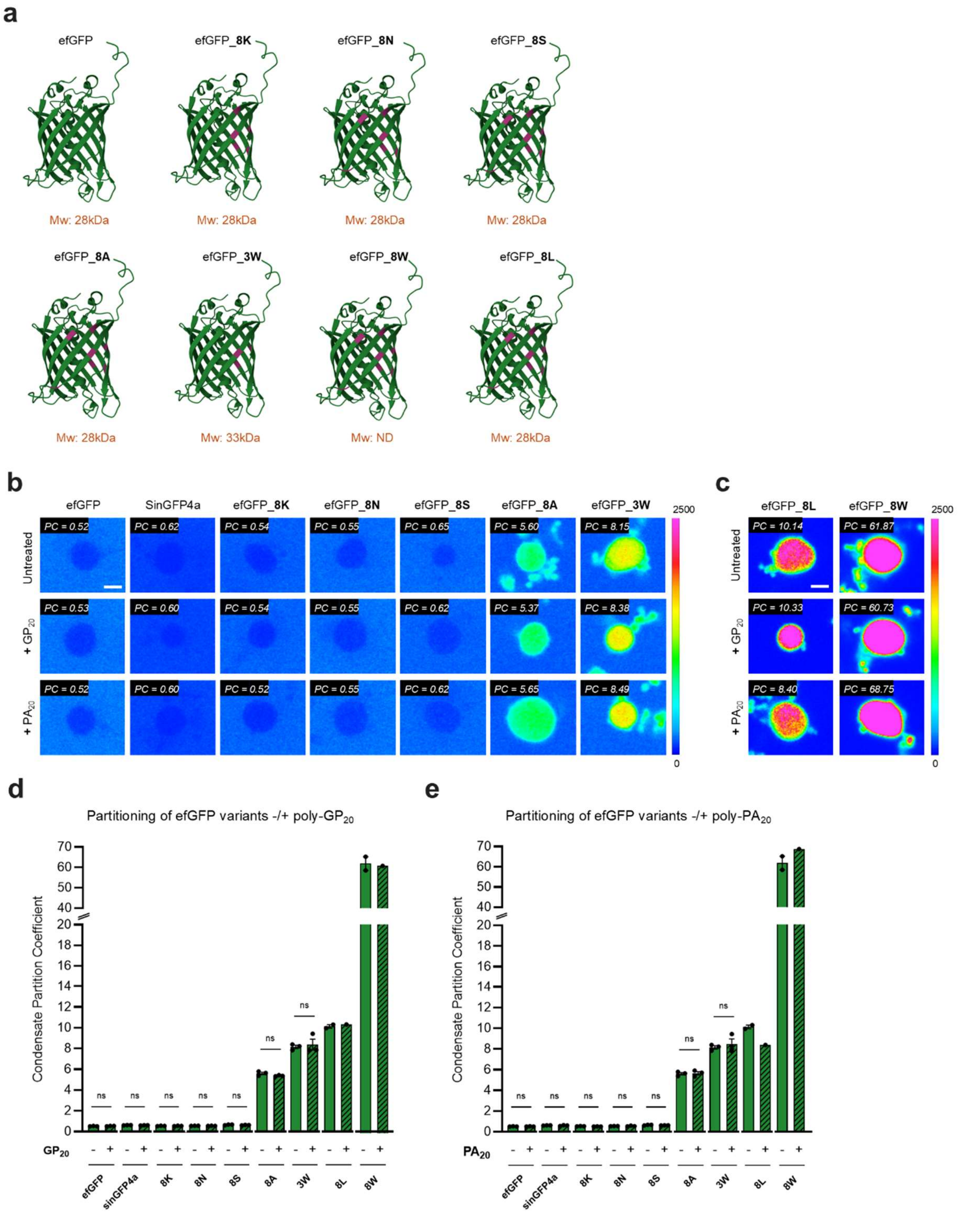


**Extended Data Figure 4 | PolyGP and polyPA do not alter FG-phase partitioning or aggregation of efGFP surface variants. a,** Eight surface residues of the efGFP scaffold were mutated (purple) to systematically alter interactions with FG domains, generating FG-philic and FG-phobic variants. AlphaFold2 predictions indicated that these mutations did not disrupt overall efGFP folding. Molecular weights from Frey *et al.*[41] indicated. **b, c, d, e,** Representative heatmap images and mean partition coefficients show that polyGP and polyPA had no detectable effect on partitioning of efGFP variants into human Nup98 FG condensates (two-tailed paired t-tests; n = 5 biological replicates, ≥5 condensates per replicate; ns, not significant). For the highly hydrophobic FG-philic variants efGFP_8L and efGFP_8W, no aggregation or altered partitioning was observed in the presence of polyGP or polyPA; (replicate numbers were insufficient to perform two-tailed paired t-tests). Scale bars = 2 µm.

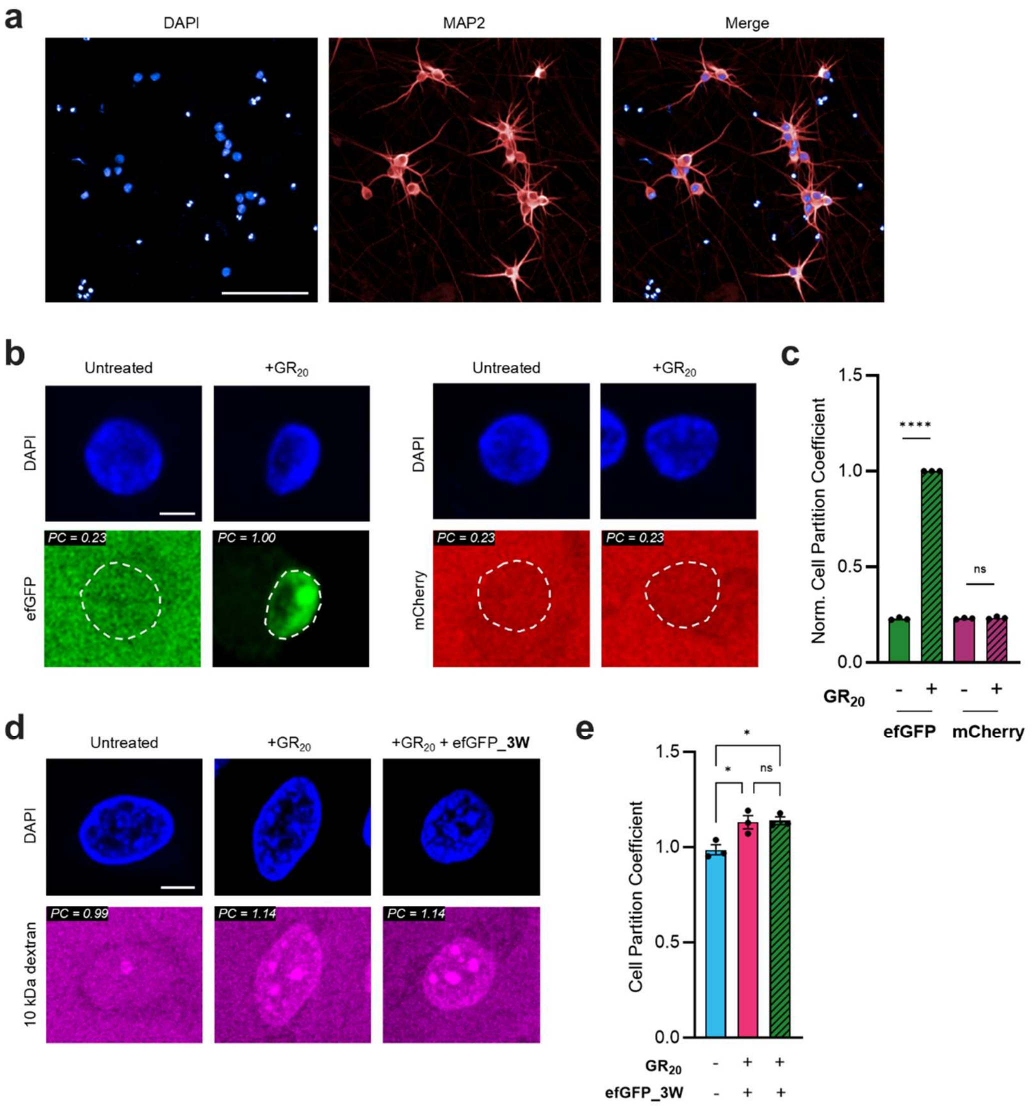


**Extended Data Figure 5 | PolyGR-dependent modulation of passive nucleocytoplasmic transport in human neurons. a,** Immunofluorescent staining showing neuronal identity by MAP2+ staining in human iPSC-derived cortical neurons (NGN2-induced) used in this study. **b,c**, Passive nuclear import of efGFP and mCherry in iPSC-derived neurons (experiments for each mutant were performed separately but assayed in parallel with efGFP, with and without $GR_{20}$, within each biological replicate; PCs are normalised to efGFP+$GR_{20}$ mean per replicate). PolyGR enhances nuclear import of wild-type efGFP (data replicated from Fig 4d,f) but has no effect on mCherry. Stats: two-tailed paired *t*-test for mCherry, one-sample *t*-test for wild-type efGFP due to normalisation; n= 3 biological replicates;. **d,e,** Passive nuclear import of 10 kDa dextran in untreated HeLa cells, or cells treated with GR20 alone or in the presence of the highly FG-philic efGFP_3W demonstrate that polyGR does not block transport, even under conditions of strong FG engagement. Stats: one-way ANOVA with post-hoc Tukey's multiple comparisons test, N=3 biological replicates. Data represent the mean ± SEM. ****p < 0.0001, *p < 0.05, ns, non-signifiant. Scale bar = 100 µm (a), 10 µm (b), 5 µm (c).

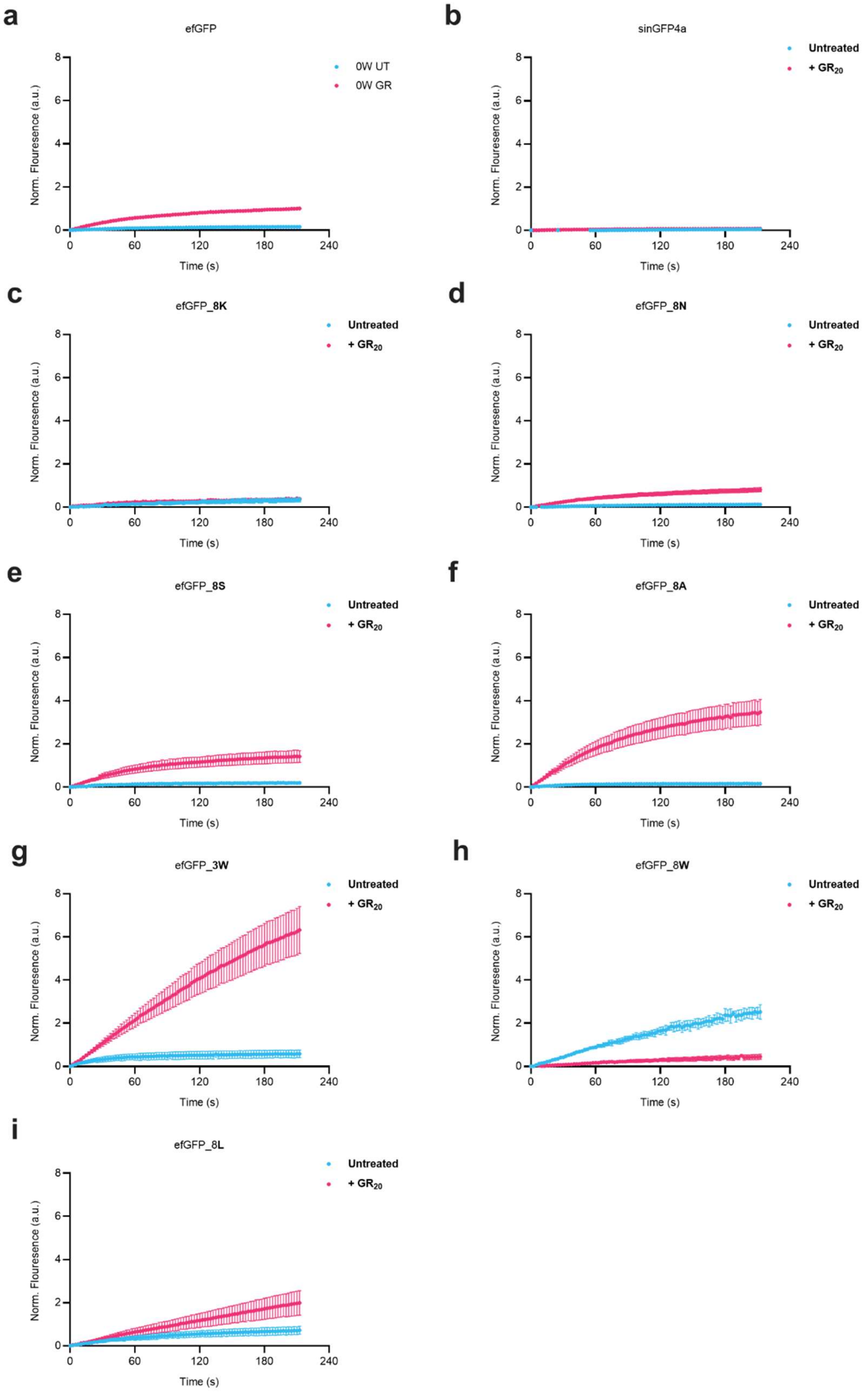


**Extended Data Figure 6 | Time courses of polyGR effects on passive nuclear import of efGFP surface variants in HeLa cells**. **a, d, e, f, g, h,** Time courses of nuclear import for efGFP surface variants in semi-permeabilised HeLa cells, either untreated or pre-treated with polyGR. PolyGR selectively altered nuclear accumulation kinetics in a surface chemistry–dependent manner: wild-type efGFP, neutral and FG-philic variants exhibited increased nuclear accumulation over time, whereas highly FG-phobic variants showed little or no change in import dynamics. **b,c,** The FG-phobic variants efGFP_8K and sinGFP4a displayed comparable nuclear import kinetics in the absence and presence of polyGR, demonstrating insensitivity to polyGR across the time course. **i,** The highly hydrophobic FG-philic variant efGFP_8W exhibited reduced nuclear accumulation over time following polyGR treatment. Together, these time-course measurements show that polyGR alters both the extent and kinetics of passive nuclear import in a client-dependent manner, reinforcing a surface chemistry–defined continuum of transport outcomes.

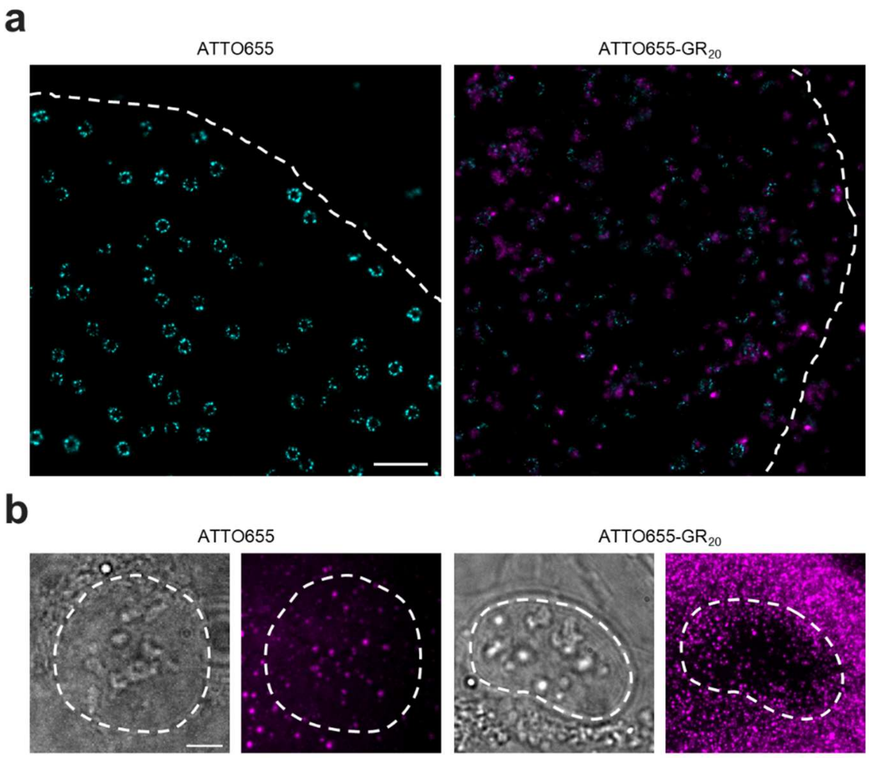


**Extended Data Figure 7 | Supporting data for super-resolution imaging of polyGR interaction with the nuclear pore complex in human cells. a,** Representative full field of view image of STORM imaging of fixed cells showing that ATTO655 dye alone does not localize to the nuclear envelope or NPCs, whereas ATTO655–$GR_{20}$ exhibits localisation to the nuclear envelope and in close proximity to NPCs. White dotted line indicates the boundary of the nuclear envelope defined by brightfield imaging. Scale bars are 500 nm. **b**, Live-cell imaging showing that both ATTO655 dye alone and ATTO655-$GR_{20}$ penetrate the plasma membrane. Scale bars are 5 μm.
