## Supplementary Table for "C9ORF72-derived polyGR polypeptides disrupt passive nucleocytoplasmic transport by tuning protein affinity for the nuclear pore barrier"

| Protein name | Construct | Source |
|---|---|---|
| Human Nup98 FG domain | $His_{14}$–TEVcs–*hs*$Nup98^{1-499}$–Cys | This study |
| efGFP* | $His_{14}$-ZZ-SUMOstar-efGFP_0W | [1] |
| efGFP_3W* | $His_{14}$-ZZ-SUMOstar-efGFP_3W | [1] |
| efGFP_8W* | $His_{14}$-ZZ-SUMOstar-efGFP_8W | [1] |
| efGFP_8L* | $His_{14}$-ZZ-SUMOstar-efGFP_8L | [1] |
| efGFP_8A* | $His_{14}$–*bd*SUMO–efGFP_8A | [1] |
| efGFP_8S* | $His_{14}$–*bd*SUMO–efGFP_8S | [1] |
| efGFP_8N* | $His_{14}$–*bd*SUMO–efGFP_8N | [1] |
| efGFP_8K* | $His_{14}$–*bd*SUMO–efGFP_8K | [1] |
| TEV protease* | MBP–TEVcs–$His_7$–TEVp(L56V,S135G,S219V) | pDZ2087[2] from Addgene |
| Ulp1Star protease* | $His_{14}$–TEVcs–Ulp1Star | [3] |
| *bd*SENP1 protease* | $His_{14}$–TEVcs–*bd*SENP1 | [3] |

**Supplementary Table 1**: **Proteins examined in this study and their corresponding bacterial expression constructs.** Entries marked with an asterisk indicate constructs from which the His tag and other tags or cleavage sites were removed by SUMO or TEV protease cleavage. TEVcs refers to the TEV cleavage site, and TEVp refers to TEV protease. The Ulp1Star protease was used to cleave SUMOstar-tagged efGFP variants, whereas bdSENP1 was used to cleave bdSUMO-tagged efGFP variants.

### Supplementary References


1. Frey, S. *et al*. Surface Properties Determining Passage Rates of Proteins through Nuclear Pores. *Cell* **174**, 202–217.e209 (2018).
2. Raran-Kurussi, S., Cherry, S., Zhang, D. & Waugh, D.S. Removal of Affinity Tags with TEV Protease, in *Heterologous Gene Expression in E.coli: Methods and Protocols*. (ed. N.A. Burgess-Brown) 221–230 (Springer, New York, NY; 2017).
3. Frey, S. & Görlich, D. A new set of highly efficient, tag-cleaving proteases for purifying recombinant proteins. *Journal of Chromatography A* **1337**, 95–105 (2014).
